# Integrative analysis of the MD Anderson Prostate Cancer Patient-Derived Xenograft Series (MDA PCa PDX)

**DOI:** 10.1101/2022.07.21.500652

**Authors:** Nicolas Anselmino, Estefania Labanca, Xiaofei Song, Jun Yang, Peter DA Shepherd, Jiabin Dong, Ritika Kundra, Nikolas Schultz, Jianhua Zhang, John C. Araujo, Ana M Aparicio, Sumit K. Subudhi, Paul G. Corn, Louis L. Pisters, John F. Ward, John W. Davis, Geraldine Gueron, Elba S. Vazquez, Christopher J Logothetis, Andrew Futreal, Patricia Troncoso, Yu Chen, Nora M Navone

## Abstract

Progress in understanding prostate cancer (PCa) metastasis and therapy resistance has been hampered by the lack of models, representative of the clinical spectrum and biologic complexity of the disease. Our laboratory is home to one of the largest worldwide repositories of PCa patient-derived xenografts (PDXs), the MDA PCa PDX series, a collection of clinically annotated PDXs reflecting the full spectrum of potentially lethal disease, that includes tumors that are not end stage and not castration-resistant PCa. We performed whole genome sequencing, targeted sequencing and RNA sequencing of 46 MDA PCa PDX models derived from biopsy and surgical specimens from 39 patients, selected in order to reflect the clinicopathological PCa subtypes (data available in cBioPortal). MDA PCa PDXs genomic characterization shows that the cohort recapitulates the mutational landscape found in PCa, highlighting the clinical relevance of these models. Interestingly and consistently with the clinic, certain models lack the typical PCa driver alterations, thus providing a suitable tool for discovery of novel drivers. Our cohort also includes PDXs derived from different areas of the same tumor and longitudinal samples, allowing to study disease heterogeneity and progression. Finally, we have developed a procedure to grow organoids from PDXs, thus providing a powerful *in vitro* platform that supports hypothesis generation, and testing of clinically relevant observations. Genomic and transcriptomic characterization of MDA PCa PDXs together with the ability to grow them as organoids for *in vitro* experimentation, provides a unique resource to address the existing clinical gap in PCa, helping to better understand mechanisms of response and resistance.

**One Sentence Summary:** MDA PCa PDX series is a dynamic resource capturing the molecular landscape of prostate cancer; a platform for discovery and personalized medicine

## Introduction

Metastatic prostate cancer (PCa) that progresses after androgen ablation therapy (i.e., castration-resistant PCa [CRPC]) remains incurable. Understanding the determinants of metastasis and therapy resistance has been hindered by the lack of models, representative of the clinical spectrum and biologic complexity of PCa. Patient-derived xenografts (PDXs) have been developed and have led to therapeutically relevant approaches (1-6). The success of these models, and recent large-scale genomics studies that have identified deregulated pathways in metastatic CRPC (mCRPC), further drives the impetus to understand and improve the utility of PDXs to bridge the gap between clinical and mechanistic observations.

The MDA PCa PDX series showcase the detailed clinical annotation of the donor tumor and the corresponding patient, enabling the understanding of cross species differences (7). We previously reported the presence of several classes of cancer mutations and gene expression alterations in these PDX models, which led to the identification of putative therapies targeting these alterations (8). We also provided evidence on the relevance of the MDA PCa PDX series in recapitulating human PCa and enabling optimization of PCa–specific marker-driven therapy (7).

Advanced PCa is characterized by somatic mutations, gene fusions and copy number variations (CNVs) (9, 10). In this work, we selected 46 MDA PCa PDX models derived from 39 patient tumors that reflect the various morphologic groups, stages, and treatment statuses of the disease. In these models, we performed whole genome sequencing (WGS), targeted sequencing and RNA sequencing (RNAseq). Genomic results of our studies have been uploaded in cBioPortal. Importantly, the molecular and morphological analyses of each PDX were all performed in representative samples of the same tumor. This design facilitates the integration of the different approaches of genomic analyses (i.e., WGS, targeted sequencing and RNAseq) with the morphologic and immunoassays results. Our genomic studies show that MDA PCa PDXs recapitulate the mutational landscape found in PCa, highlighting the clinical relevance of the cohort. We also present evidence that PDXs can be propagated as organoids to perform drug testing or functional studies.

## Results

The MDA PCa PDX series is a collection of clinically annotated PCa PDXs representing the whole range of potentially lethal disease, namely therapy-naïve and therapy-resistant PCa encompassing clinical (classic and aggressive) and morphological (e.g., adenocarcinomas (Ad), neuroendocrine (NE) carcinoma) variants derived from both primary sites and metastases, reflecting the therapy progression of the tumor donor, which is monitored by genitourinary oncologists (11, 12). This collection was created by developing a strategy to establish PCa PDXs from tumor specimens obtained from patients with PCa undergoing radical prostatectomy, cystoprostatectomy/pelvic exenteration, or resection/biopsy analysis of metastatic lesions. By generating PDXs from different areas of the same tumor, we have developed models of PCa heterogeneity. Compared with other banks, our Program collects samples that are not end stage disease. Our dynamic series also includes models taken at different time points, enabling longitudinal studies. These PDXs are submitted to rigorous quality control.

As the MDA PCa PDX program is constantly accruing samples for PDX development, it captures the evolving molecular landscape of PCa progressing under conventional and novel therapies (1^st^ and 2^nd^ generation androgen deprivation therapies, chemotherapy, and targeted therapies). To date, we have developed two cell lines (MDA PCa 2a and MDA PCa 2b) and 139 PDXs derived from 86 PCa patients. The racial distribution of patient donors of the selected PDXs for these studies reflects the patient population treated at our institution (32 Caucasian, 5 African American, and 2 Hispanic).

We performed WGS, targeted sequencing for 263 gene mutations implicated in the pathogenesis of solid cancers (T200.1 panel) (Table S1), and RNAseq of 46 MDA PCa PDX models derived from 39 patient tumors. These 46 PDXs were selected to reflect the various morphologic groups, stages, and treatment statuses of the disease (Fig 1). Table 1 outlines the clinical and morphological features of these PDXs.

**Figure 1.**
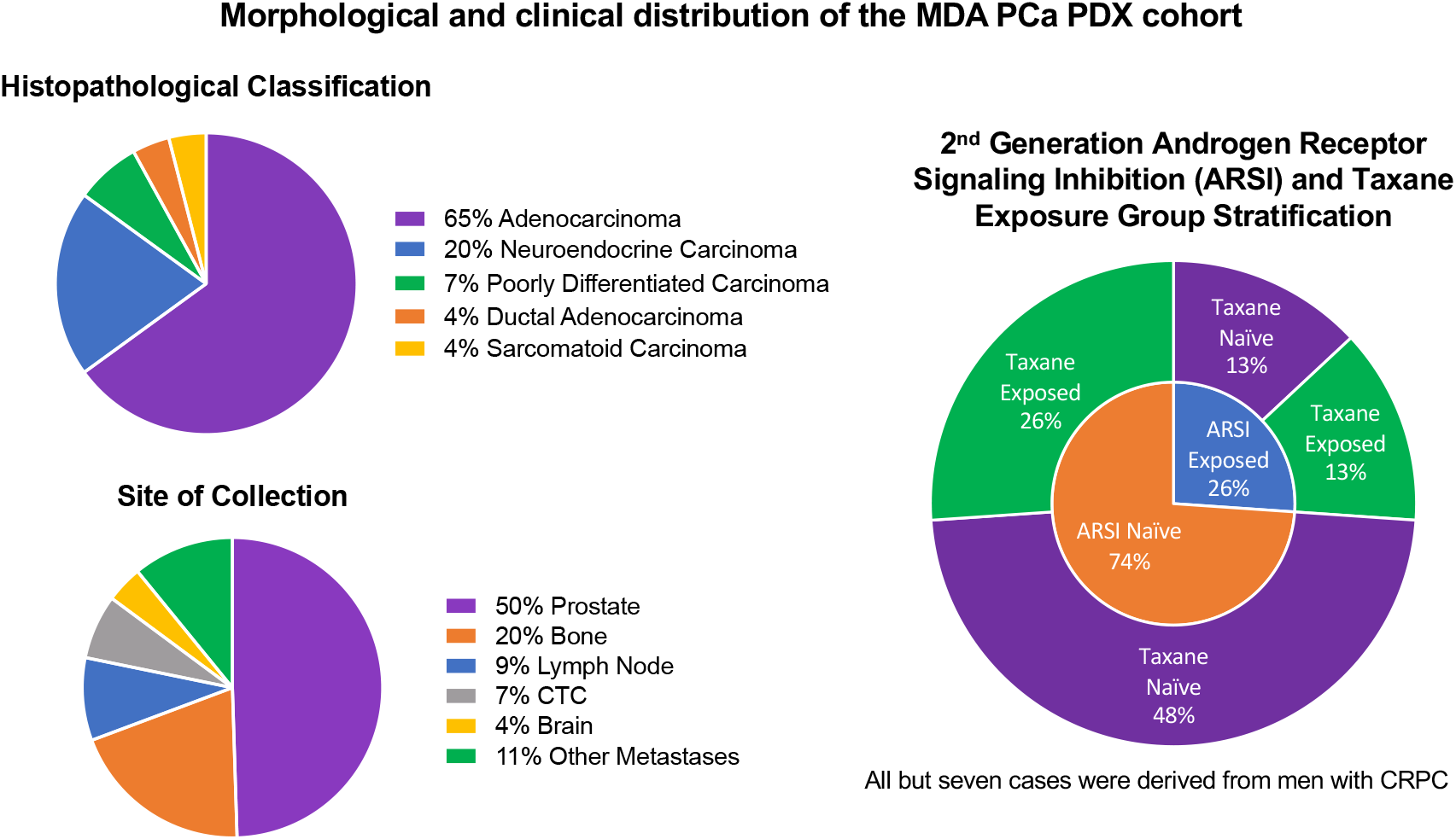
Morphological and clinical distribution of the MDA PCa PDX (patient-derived xenograft) cohort. Previous treatment, morphological classification and site of tumor collection of the 46 PDXs (from 39 prostate cancer patients) used for genomic analysis (whole genome sequencing (low-pass) (WGS), targeted sequencing (T200.1 panel; 263 genes) and RNA sequencing). CTC: circulating tumor cells; CRPC: castration-resistant prostate cancer.

**Table 1.** Clinical and morphological features of 46 MPA PCa PDXs.

| Patient ID | PDX name is "MDA PCa+the alphanumeric string below" | Race | Treatment Status | Last Treatment Category | Prior Treatments | Primary, Bone or other metastases | Tumor Site | Morphology of PDX |
| --- | --- | --- | --- | --- | --- | --- | --- | --- |
| 1 | 117-9 | Hispanic | mCRPC | Chemotherapy | 1st Generation ADT | Primary | Prostate | Adenocarcinoma |
| 2 | 118-B | White | mCRPC | Chemotherapy | 1st Generation ADT | Bone | Bone | Adenocarcinoma |
| 3 | 133-4 | White | mCRPC | Chemotherapy | 1st Generation ADT | Bone | Bone | Adenocarcinoma |
| 4 | 144-13 | White | mCRPC | Chemotherapy | 1st Generation ADT | Primary | Prostate | Neuroendocrine Carcinoma |
| 4 | 144-4 | White | mCRPC | Chemotherapy | 1st Generation ADT | Primary | *Bladder | Neuroendocrine Carcinoma |
| 5 | 146-10 | White | mCRPC | Chemotherapy | 1st Generation ADT | Primary | *Bladder | Neuroendocrine Carcinoma |
| 5 | 146-12 | White | mCRPC | Chemotherapy | 1st Generation ADT | Primary | *Bladder | Adenocarcinoma |
| 6 | 150-7 | White | mCRPC | Chemotherapy | 1st Generation ADT | Bone | Bone | Neuroendocrine Carcinoma |
| 7 | 152-1 | White | mCRPC | 1st Generation ADT |  | Other Metastasis | Brain | Ductal Adenocarcinoma |
| 8 | 153-7 | White | mCRPC | 1st Generation ADT |  | Other Metastasis | Thyroid | Adenocarcinoma |
| 9 | 155-2 | White | mCRPC | Chemotherapy | 1st Generation ADT | Primary | *Bladder | Neuroendocrine Carcinoma |
| 10 | 163-A | White | mCRPC | 1st Generation ADT |  | Primary | Prostate | Adenocarcinoma |
| 11 | 166-1 | White | mCRPC | Chemotherapy | 1st Generation ADT | Primary | *Bladder | Adenocarcinoma |
| 12 | 173-2 | White | Naïve | Naïve |  | Primary | Prostate | Adenocarcinoma |
| 13 | 174-6 | Black | mCRPC | Chemotherapy | 1st Generation ADT | Primary | Prostate | Adenocarcinoma |
| 14 | 175-6 | Hispanic | mCRPC | Chemotherapy | 1st Generation ADT | Other Metastasis | Testis | Adenocarcinoma |
| 15 | 177-B | White | mCRPC | 1st Generation ADT |  | Primary | Prostate | Poorly Differentiated Carcinoma |
| 16 | 178-11 | White | NADT | 1st Generation ADT |  | Primary | Prostate | Adenocarcinoma |
| 17 | 180-30 | White | mCRPC | Chemotherapy | 1st Generation ADT | Primary | Prostate | Adenocarcinoma |
| 18 | 183-A | White | Naïve | Naïve |  | Bone | Bone | Adenocarcinoma |
| 15 | 189-1 | White | mCRPC | Chemotherapy | 1st Generation ADT | Primary | Prostate | Adenocarcinoma |
| 19 | 200-1 | White | mCRPC | 1st Generation ADT |  | Bone | Bone | Adenocarcinoma |
| 18 | 203-A | White | mCRPC | 1st Generation ADT |  | Bone | Bone | Adenocarcinoma |
| 20 | 250-15 | White | NADT | 1st Generation ADT |  | Other Metastasis | Adrenal Gland | Ductal Adenocarcinoma |
| 21 | 265-8 | White | mCRPC | Chemotherapy | 2nd Generation ADT | Primary | Prostate | Adenocarcinoma |
| 22 | 266-A | White | mCRPC | Chemotherapy | 1st Generation ADT | Other Metastasis | Serous Cavity | Adenocarcinoma |
| 23 | 270-A | White | NADT | 1st Generation ADT |  | Bone | Bone | Adenocarcinoma |
| 24 | 273-A | White | mCRPC | 2nd Generation ADT |  | Other Metastasis | Lymph Node | Neuroendocrine Carcinoma |
| 25 | 274-2 | White | mCRPC | Chemotherapy | 2nd Generation ADT | Bone | Bone | Adenocarcinoma |
| 26 | 277-6 | White | mCRPC | 2nd Generation ADT |  | Other Metastasis | Lymph Node | Neuroendocrine Carcinoma |
| 27 | 280-9 | White | Naïve | Naïve |  | Other Metastasis | Lymph Node | Adenocarcinoma |
| 28 | 285-3 | White | mCRPC | 2nd Generation ADT |  | Primary | *Bladder | Adenocarcinoma |
| 27 | 306-14 | White | mCRPC | 1st Generation ADT |  | Primary | Prostate | Adenocarcinoma |
| 29 | 312-7 | White | mCRPC | Chemotherapy | 1st Generation ADT | Primary | Prostate | Neuroendocrine Carcinoma |
| 30 | 315-8 | White | mCRPC | 2nd Generation ADT |  | Primary | Prostate | Poorly Differentiated Carcinoma |
| 31 | 316-1 | White | mCRPC | 2nd Generation ADT |  | Bone | Bone | Adenocarcinoma |
| 31 | 316-2 | White | mCRPC | 2nd Generation ADT |  | Other Metastasis | CTC | Adenocarcinoma |
| 32 | 320-1 | White | mCRPC | 1st Generation ADT |  | Other Metastasis | CTC | Adenocarcinoma |
| 33 | 322-2 | Black | NADT | 1st Generation ADT |  | Primary | Prostate | Adenocarcinoma |
| 34 | 327-2 | White | mCRPC | Chemotherapy | 2nd Generation ADT | Other Metastasis | CTC | Adenocarcinoma |
| 35 | 328-5 | White | CRPC | 1st Generation ADT |  | Primary | *Bladder | Neuroendocrine Carcinoma |
| 36 | 337-A | Black | mCRPC | 1st Generation ADT |  | Other Metastasis | Liver | Poorly Differentiated Carcinoma |
| 37 | 342-B | White | mCRPC | Chemotherapy | 1st Generation ADT | Primary | Prostate | Sarcomatoid Carcinoma |
| 38 | 352-8 | Black | mCRPC | 2nd Generation ADT |  | Other Metastasis | Lymph Node | Adenocarcinoma |
| 37 | 355-9 | White | mCRPC | Chemotherapy | 2nd Generation ADT | Primary | *Bladder | Sarcomatoid Carcinoma |
| 39 | 377-1 | Black | mCRPC | Chemotherapy | 2nd Generation ADT | Other Metastasis | Brain | Adenocarcinoma |

NADT, On Neoadjuvant androgen deprivation therapy; \*, Direct extension of a prostate cancer; CTC, circulating tumor cells
PDXs with same Patient ID are derived from diferent areas of the same tumor at the same time

Morphologically, the more represented groups in the cohort are adenocarcinoma (Ad) and neuroendocrine carcinoma (NEC), defined morphologically and by negative staining for androgen receptor (AR) and positive staining for neuroendocrine markers (Chromogranin, synaptophysin, or CD56/NCAM1). When we assessed the driver mutation frequency of T200.1 genes, we found *RB1* as the only one with a significant difference between these groups, displaying a higher frequency in NEC (P=0.047) accompanied by decreased expression (P=0.005) (13-15).

### Copy number variations

We have analyzed WGS results to identify CNVs that affect cancer genes and integrated these data with mRNA expression levels to determine the driver genes that could be regulated by these alterations. In general, we found that heterozygous deletion or amplification of specific genes would not result in altered expression levels of the gene as assessed by RNAseq. However, most homozygous deletions resulted in *null* expression of the gene (Fig 2 and see cBioportal).

**Figure 2.**
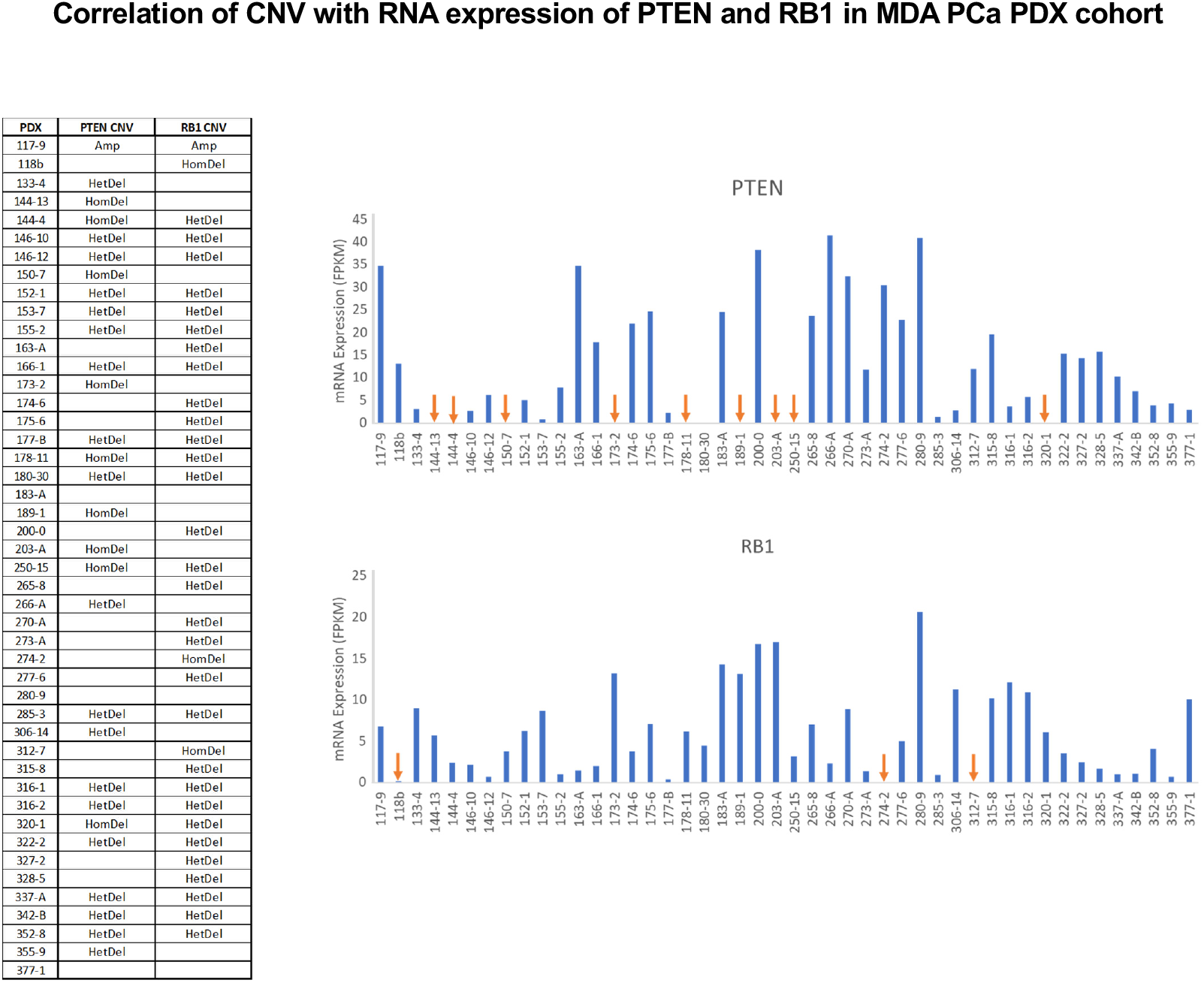
Correlation of copy number variation (CNV) with RNA expression of *PTEN* and *RB1* in MDA PCa PDX cohort. CNV of Phosphatase And Tensin Homolog (*PTEN)* and Retinoblastoma 1 (*RB1)*, based on whole genome sequencing (WGS) data (left) and RNA expression levels (right). Copy number status was categorized using log2.mean cutoffs: ≥ 0.4, Amplification (Amp); −2 to −0.4, Heterozygous Deletion (HetDel); ≤ −2 Homozygous Deletion (HomDel). Blank spaces in the table correspond to log2.mean values between −0.4 to 0.4 (absence of CNV). Orange arrows indicate models with HomDel. FPKM: fragments per kilobase of transcript per million mapped reads.

### Mutational landscape of known PCa associated genes

When we looked at SNPs/indels and copy number changes identified by targeted sequencing and WGS, we found that most pathways frequently altered in PCa (namely androgen receptor (AR), Cell cycle, DNA repair, receptor tyrosine kinase (RTK)-RAS, phosphatidylinositol 3 kinase (PI3K) pathway, Wnt/β-catenin and Chromatin modifiers) as well as copy number gains and losses (e.g., 8q and 8p, respectively) frequently found in PCa (16), were also observed in our cohort of PDXs (Fig 3). This reveals that critical aspects of clinical PCa are reflected in the PDXs.

**Figure 3.**
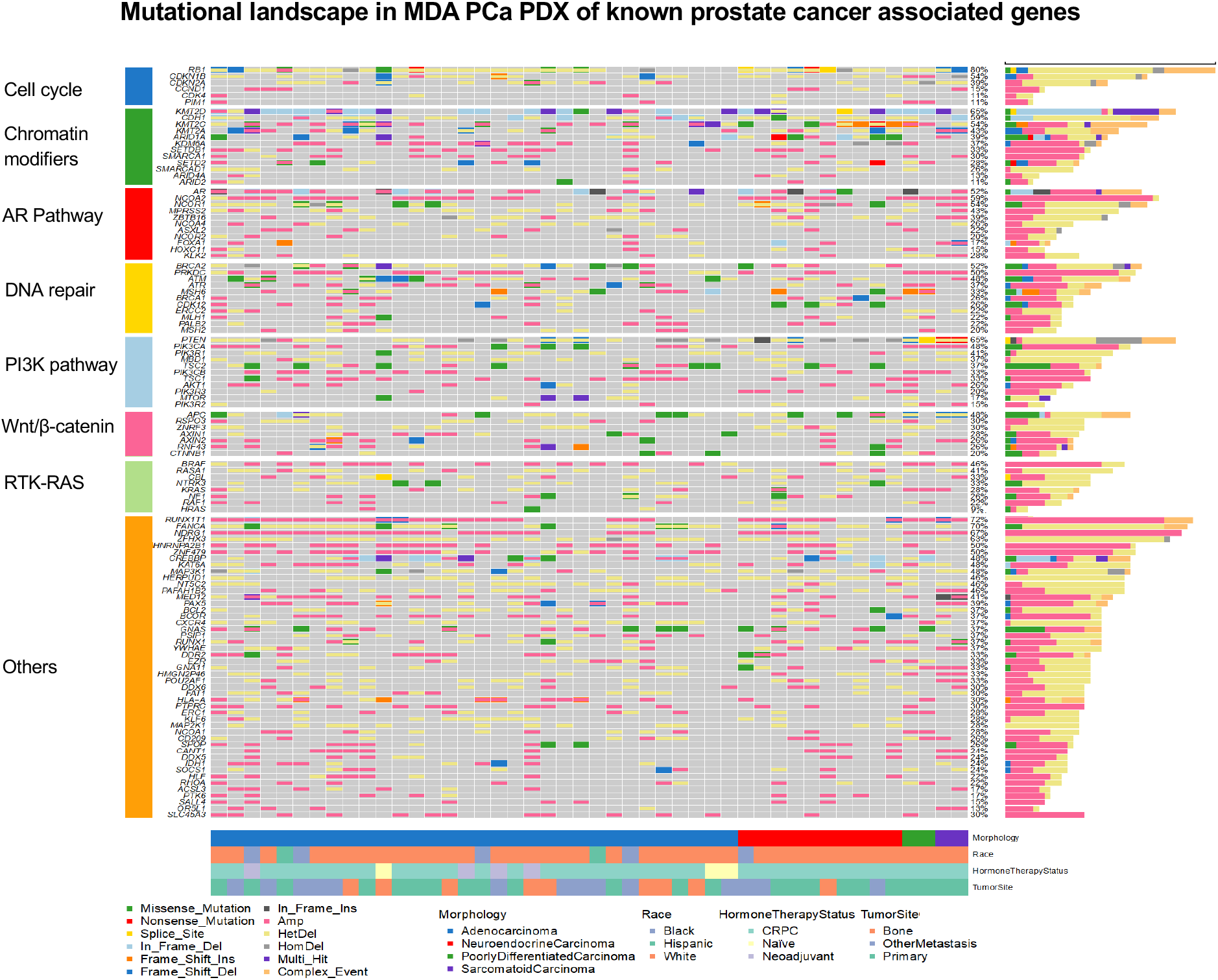
Mutational landscape in MDA PCa PDX of known prostate cancer associated genes. Single nucleotide polymorphism (SNP)/Insertion & deletion (Indel) and copy number variation (CNV) of known prostate cancer associated genes identified by targeted sequencing (T200.1 panel) and whole genome sequencing (WGS), respectively, in the 46 MDA PCa PDXs studied. Genes are grouped based on the pathways they are involved in, depicted on the left. AR: androgen receptor; PI3K: phosphatidylinositol 3 kinase; RTK-RAS: receptor tyrosine kinase-RAS; Ins: insertions; Del: deletions; Amp: amplification; HomDel: homozygous deletion; HetDel: heterozygous deletion; CRPC: castration-resistant prostate cancer.

### Fusions

Several fusions were identified in our PDX cohort at both DNA and RNA levels. We focused on fusions previously identified in human PCa (12, 17-20). Herein we report fusions that were detected at both DNA and RNA levels (of note, the breakpoint identified at DNA or RNA level may be slightly different) (Fig 4 and Table S2).

**Figure 4.**
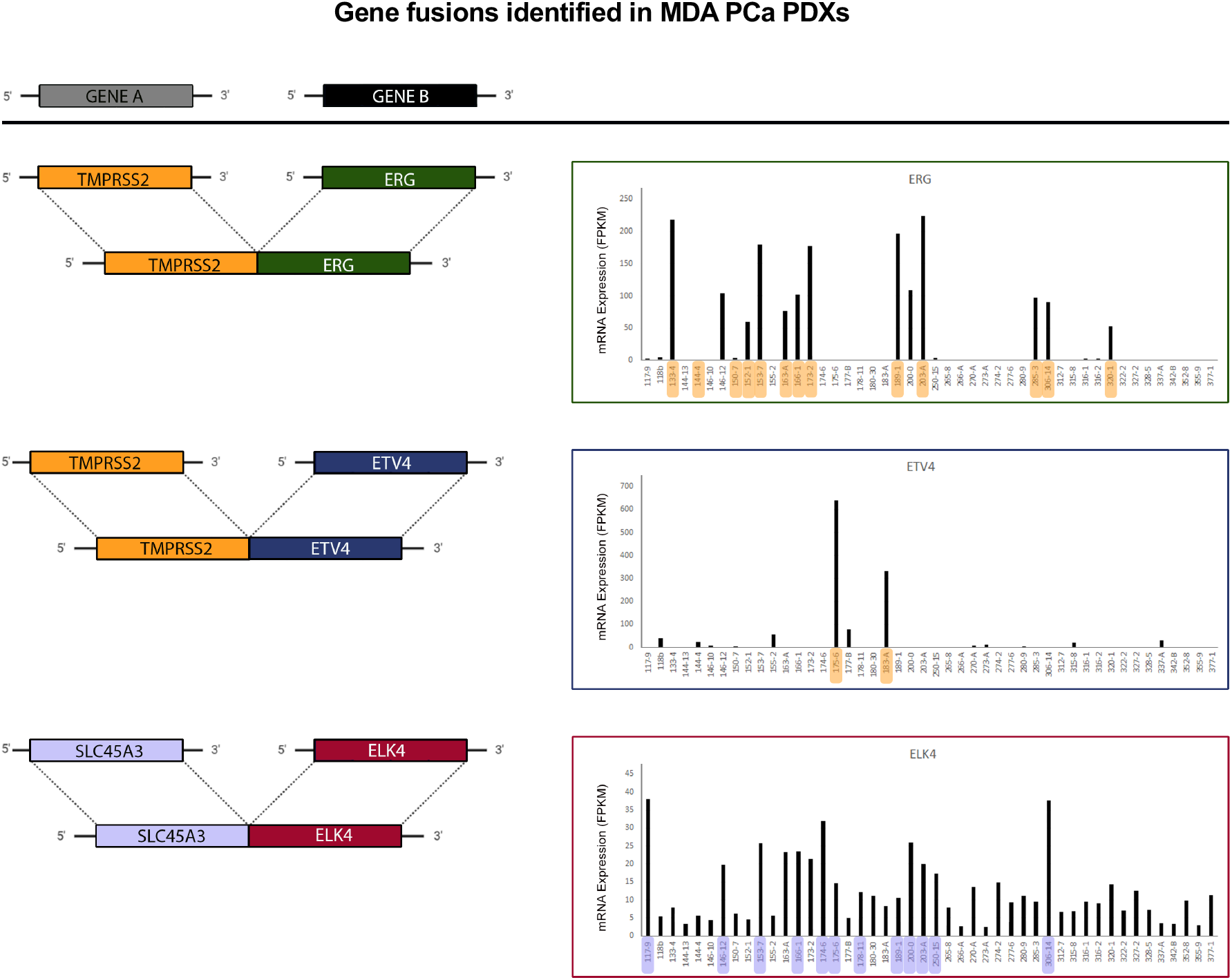
Gene fusions identified in MDA PCa PDXs. Representation of fusions identified (left) associated with high RNA expression levels of gene B (right) in the MDA PCa PDX cohort studied. Models in which fusions were detected are highlighted in orange for TMPRSS2 and in light violet for SLC45A3. TMPRSS2: transmembrane serine protease 2; ERG: ETS-related gene; ETV4: ETS variant transcription factor 4; ELK4: ETS transcription factor ELK4; SLC45A3: Solute Carrier Family 45 Member 3; FPKM: fragments per kilobase of transcript per million mapped reads.

We found that transmembrane serine protease 2-ETS-related gene (TMPRSS2-ERG) fusion correlated with increased expression in ERG in MDA PCa 133-4, 152-1, 153-7, 163-A, 166-1, 173-2, 189-1, 203-A, 285-3, 306-14, 320-1 (Fig 4). Although the same fusion was found in other cases, it did not correlate with an increased expression of ERG (Fig 4). In many of these cases, this could be explained by the low expression of AR, a known positive regulator of TMPRSS2. ERG is a frequent fusion partner of TMPRSS2, yet fusions with other members of the ETS family are also found in Pca (20). In our cohort, we also identified TMPRSS2-ETV4 fusion with increased expression in ETV4 in MDA PCa 175-6 and MDA PCa 183-A (Fig 4). ETV1, other member of the ETS family, was found in four other fusions (ETV1-FOXA1, FOXA1-ETV1, ACSL3-ETV1 and ETV1-ACSL3), but we did not detect changes in the expression of these genes. The biological consequence of these fusions remains to be elucidated. We also showcase the SLC45A3-ELK4 fusion, although not detected at the genome level but known as a posttranscriptional fusion, reported in PCa (21, 22). SLC45A3-ELK4 was found on MDA PCa 117-9, 174-6 and 306-14 with increased expression of ELK4 (Fig 4).

### PCa drivers

We subsequently looked at the most frequently altered genes in PCa (*AR*, Retinoblastoma 1 (*RB1), TP53* and Phosphatase And Tensin Homolog (*PTEN*). We found that most PDXs present oncogenic molecular alterations (i.e., mutations, homozygous deletions) in one or more of these genes. Fig 5A illustrates gene alterations in each PDX and its RNA expression. Some cases present low expression of *RB1* (MDA PCa 155-2, 273-A) or *PTEN* (MDA PCa 180-30, 306-14) with concomitant heterozygous deletion. Of note, PTEN has low RNA expression in MDA PCa 377-1; however, we only found homozygous deletion of the segment encompassing the gene, but the gene itself was not lost (Fig 5A). This may suggest that the lost regions may have important regulatory roles on gene expression. Interestingly, AR has high expression in MDA PCa 265-8 in the absence of alterations. Nevertheless, 13% PDXs lack alterations in common PCa drivers (Fig 5A, PDXs highlighted in orange).

**Figure 5.**
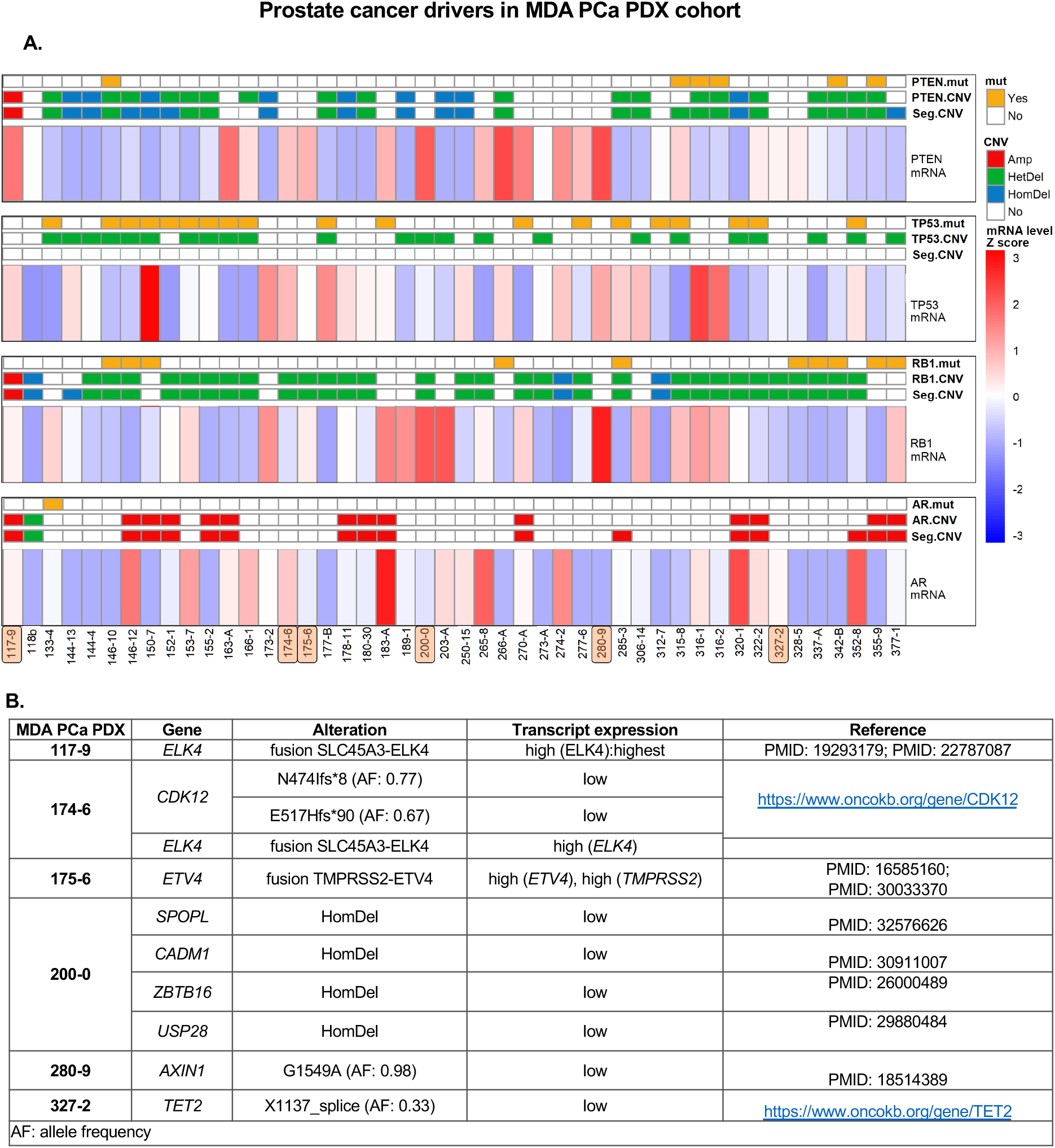
Canonical and non-canonical prostate cancer drivers in MDA PCa PDX cohort. **A**. Mutation (mut), copy number variations (CNV) and mRNA level status of genes frequently altered in prostate cancer. Highlighted in orange are the models with no relevant alterations in the genes analyzed. **B**. Putative non-canonical prostate cancer drivers in models lacking canonical alterations. AR: androgen receptor; PTEN: phosphatase and tensin homolog; Amp: amplification; HomDel: homozygous deletion; HetDel: heterozygous deletion; Seg: segment.

Next, we analyzed which genes may be driving PCa progression in this subgroup. We looked at other reported molecular alterations in PCa and in cancer in general. Fig 5B outlines our findings of putative driver genes of these PDXs. Mainly, we found alterations in genes from the ETS family, a field of study that gained relevance in the last years (19, 23). Of note, 280-9 has *null* AR by immunohistochemistry (IHC) and very low RNA levels, and although this is not *per se* a molecular alteration, AR *null* has a profound effect on the morphology and biology of PCa. Interestingly, MDA PCa 175-6 has the highest expression of BRAF among all PDXs but the gene does not have the known oncogenic mutation (V600E). It remains to validate the oncogenic role of these alterations or identify others (Fig 5B).

### AR

In our PDX cohort, only MDA PCa 133-4 has a known oncogenic mutation on AR (H875Y). When we look at CNV involving AR, we found that CNV is not always associated with altered AR expression as assessed by RNAseq (Fig 6). In fact, three cases that had AR amplifications (MDA PCa 150-7, 155-2, 355-9), had *null* expression of the receptor. On the other hand, MDA PCa 118b shows heterozygous deletion and lacks AR expression, indicating the relevance of other mechanisms in regulating AR.

**Figure 6.**
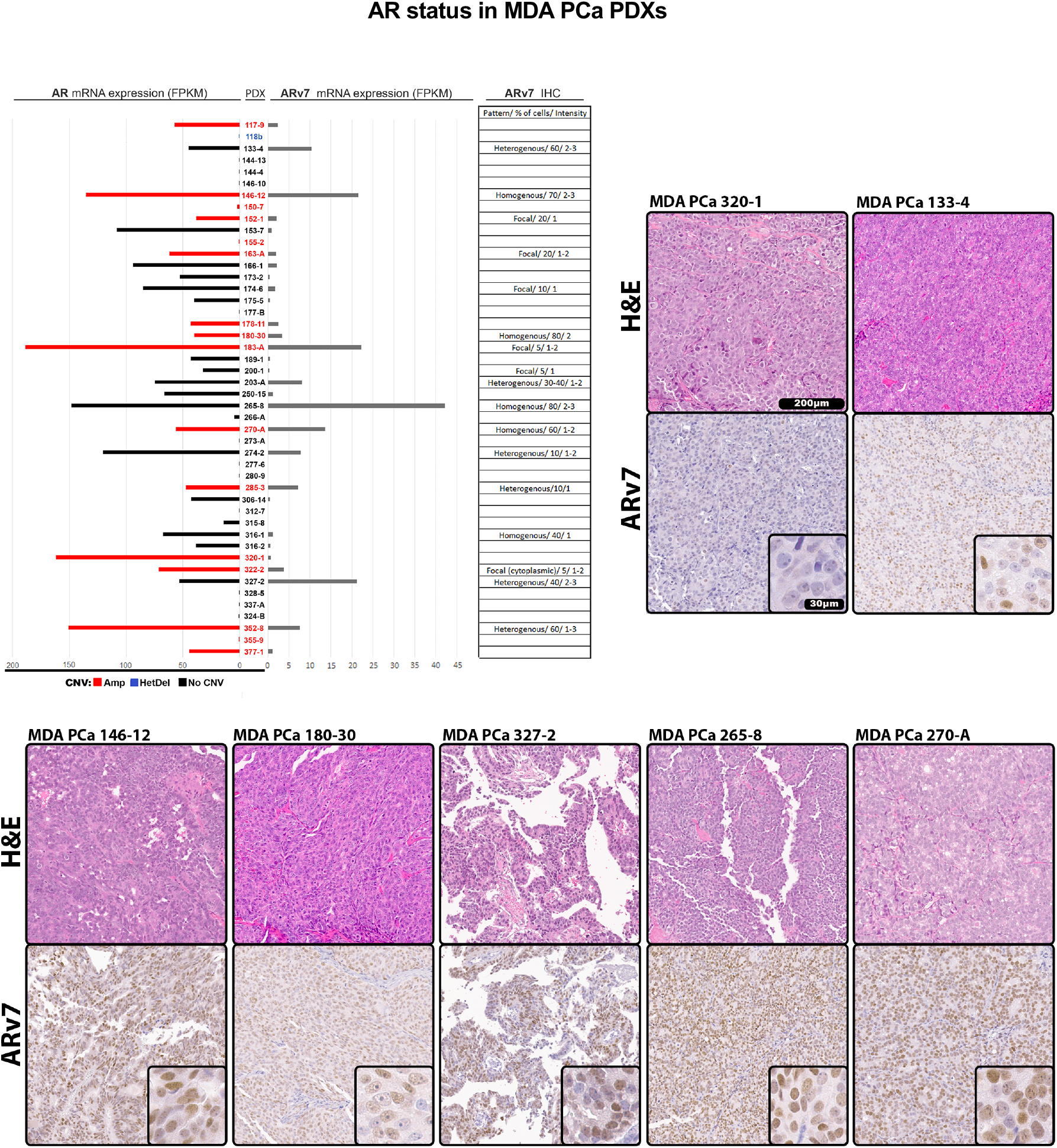
Androgen receptor (AR) status in MDA PCa PDXs. Expression of AR and AR splice variant 7 (ARv7) evaluated from RNAseq data in MDA PCa PDXs. AR bars in the chart are depicted in red for AR amplifications (Amp), blue for heterozygous deletion (HetDel) or black for no copy number variation (no CNV). Table and representative images show immunohistochemistry (IHC) for ARv7 performed in samples from the 46 MDA PCa PDX models. Blank spaces in table correspond to negative staining. Staining was evaluated by semi-quantitative analysis of pattern, percentage of cells and intensity on a scale of 1 to 3. FPKM: fragments per kilobase of transcript per million mapped reads.

Upon further analysis of *AR* expression, we then looked for the presence of the best characterized and most prevalent AR splice variant, *ARv7*. As a constitutively active variant, ARv7 is a known PCa driver. Using Integrative Genomics Viewer (IGV) software, we analyzed RNAseq data for the inclusion or not of cryptic exon corresponding to *ARv7* (Fig 6). We then confirmed the presence of ARv7 by IHC (Fig 6). We detected different *ARv7* expression levels and found that MDA PCa 146-12, 265-8, 270-8, 327-2 are the ones with highest expression of this variant. Of note, 327-2 was grouped among the 13% PDXs that did not have oncogenic alterations in *PTEN, TP53, AR*, and *RB1* (Fig 5A). It remains unknown whether ARv7 could be a PCa driver by itself or with other putative non-canonical drivers (e.g. *TET2)* (Fig 5B).

### Genomic analysis of two areas of the same tumor

Some PDXs selected for this genomic analysis are derived from different areas of the same tumor reflecting its heterogeneity (Table 1, rows with the same ID and surrounded by double borders, Table 2), namely MDA PCa 144-13 and 144-4 (Patient ID 4), MDA PCa 146-10 and 146-12 (Patient ID 5). Two other PDXs were developed from a bone metastasis and circulating tumor cells (CTC) obtained at the same time from the same patient (MDA PCa 316-1 and 316-2 (patient ID 31)).

**Table 2.**
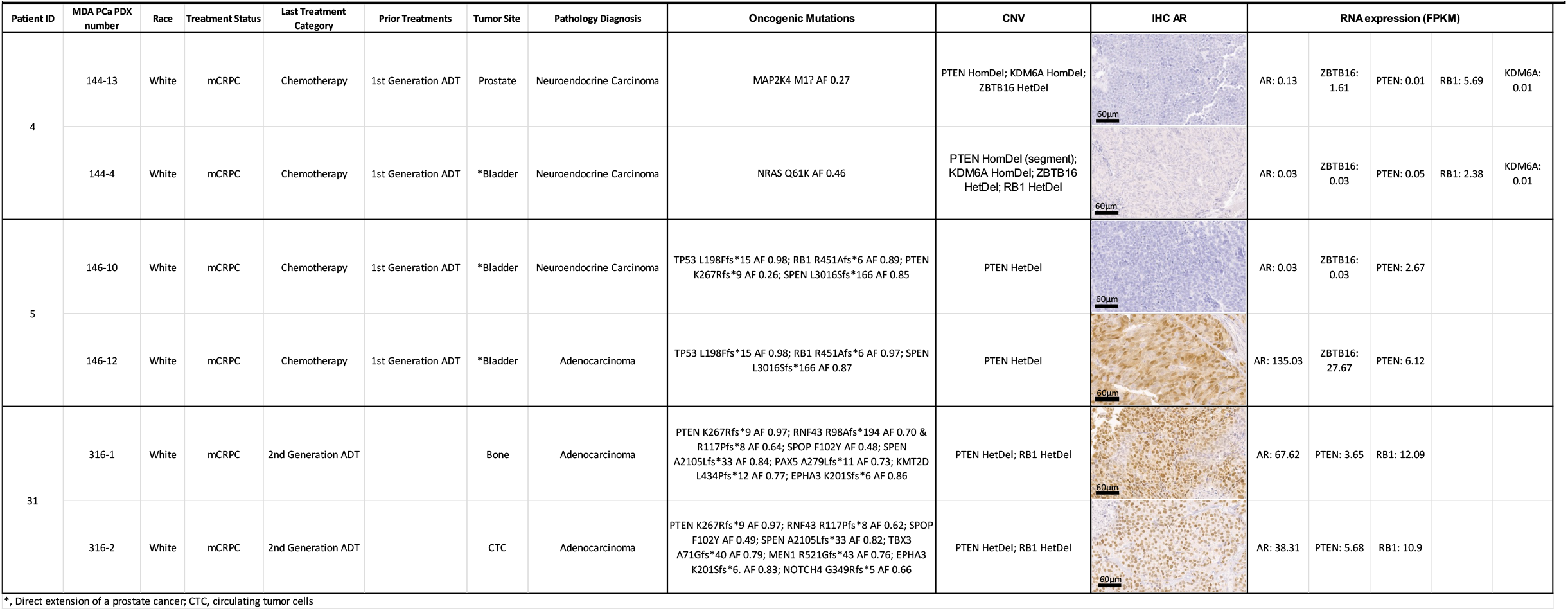
Clinical, morphological and molecular features of MDA PCa PDXs pairs derived from the same tumor.

| Patient ID | MDA PCa PDX number | Race | Treatment Status | Last Treatment Category | Prior Treatments | Tumor Site | Pathology Diagnosis | Oncogenic Mutations | CNV | IHC AR | RNA expression (FPKM) |  |  |  |  |
| --- | --- | --- | --- | --- | --- | --- | --- | --- | --- | --- | --- | --- | --- | --- | --- |
| 4 | 144-13 | White | mCRPC | Chemotherapy | 1st Generation ADT | Prostate | Neuroendocrine Carcinoma | MAP2K4 M1? AF 0.27 | PTEN HomDel; KDM6A HomDel; ZBTB16 HetDel |  | AR: 0.13 | ZBTB16: 1.61 | PTEN: 0.01 | RB1: 5.69 | KDM6A: 0.01 |
|  | 144-4 | White | mCRPC | Chemotherapy | 1st Generation ADT | *Bladder | Neuroendocrine Carcinoma | NRAS Q61K AF 0.46 | PTEN HomDel (segment); KDM6A HomDel; ZBTB16 HetDel; RB1 HetDel |  | AR: 0.03 | ZBTB16: 0.03 | PTEN: 0.05 | RB1: 2.38 | KDM6A: 0.01 |
| 5 | 146-10 | White | mCRPC | Chemotherapy | 1st Generation ADT | *Bladder | Neuroendocrine Carcinoma | TP53 L198Ffs*15 AF 0.98; RB1 R451Afs*6 AF 0.89; PTEN K267Rfs*9 AF 0.26; SPEN L3016Sfs*166 AF 0.85 | PTEN HetDel |  | AR: 0.03 | ZBTB16: 0.03 | PTEN: 2.67 |  |  |
|  | 146-12 | White | mCRPC | Chemotherapy | 1st Generation ADT | *Bladder | Adenocarcinoma | TP53 L198Ffs*15 AF 0.98; RB1 R451Afs*6 AF 0.97; SPEN L3016Sfs*166 AF 0.87 | PTEN HetDel |  | AR: 135.03 | ZBTB16: 27.67 | PTEN: 6.12 |  |  |
| 31 | 316-1 | White | mCRPC | 2nd Generation ADT |  | Bone | Adenocarcinoma | PTEN K267Rfs*9 AF 0.97; RNF43 R98Afs*194 AF 0.70 & R117Pfs*8 AF 0.64; SPOP F102Y AF 0.48; SPEN A2105Lfs*33 AF 0.84; PAX5 A279Lfs*11 AF 0.73; KMT2D L434Pfs*12 AF 0.77; EPHA3 K201Sfs*6 AF 0.86 | PTEN HetDel; RB1 HetDel |  | AR: 67.62 | PTEN: 3.65 | RB1: 12.09 |  |  |
|  | 316-2 | White | mCRPC | 2nd Generation ADT |  | CTC | Adenocarcinoma | PTEN K267Rfs*9 AF 0.97; RNF43 R117Pfs*8 AF 0.62; SPOP F102Y AF 0.49; SPEN A2105Lfs*33 AF 0.82; TBX3 A71Gfs*40 AF 0.79; MEN1 R521Gfs*43 AF 0.76; EPHA3 K201Sfs*6 AF 0.83; NOTCH4 G349Rfs*5 AF 0.66 | PTEN HetDel; RB1 HetDel |  | AR: 38.31 | PTEN: 5.68 | RB1: 10.9 |  |  |

\*, Direct extension of a prostate cancer; CTC, circulating tumor cells

MDA PCa 144-13 and 144-4 were derived from different areas of a neuroendocrine carcinoma in a primary PCa, both having *null* AR expression, as expected. They share genomic alterations in genes known to be drivers of PCa (homozygous deletion in *PTEN* and *KDM6A* and heterozygous deletion of *ZBTB16*) (Table 2, Fig S2) in line with *null* or almost *null* expression. Only MDA PCa 144-4 has heterozygous deletion of *RB1*, and that is accompanied with low expression (Table 2, Fig S2). Interestingly, MDA PCa 144-4 has a known oncogenic mutation in *NRAS* (Q61K) (Fig 7, Table 2, see cBioPortal), that has been seldom implicated in PCa pathogenesis (24).

**Figure 7.**
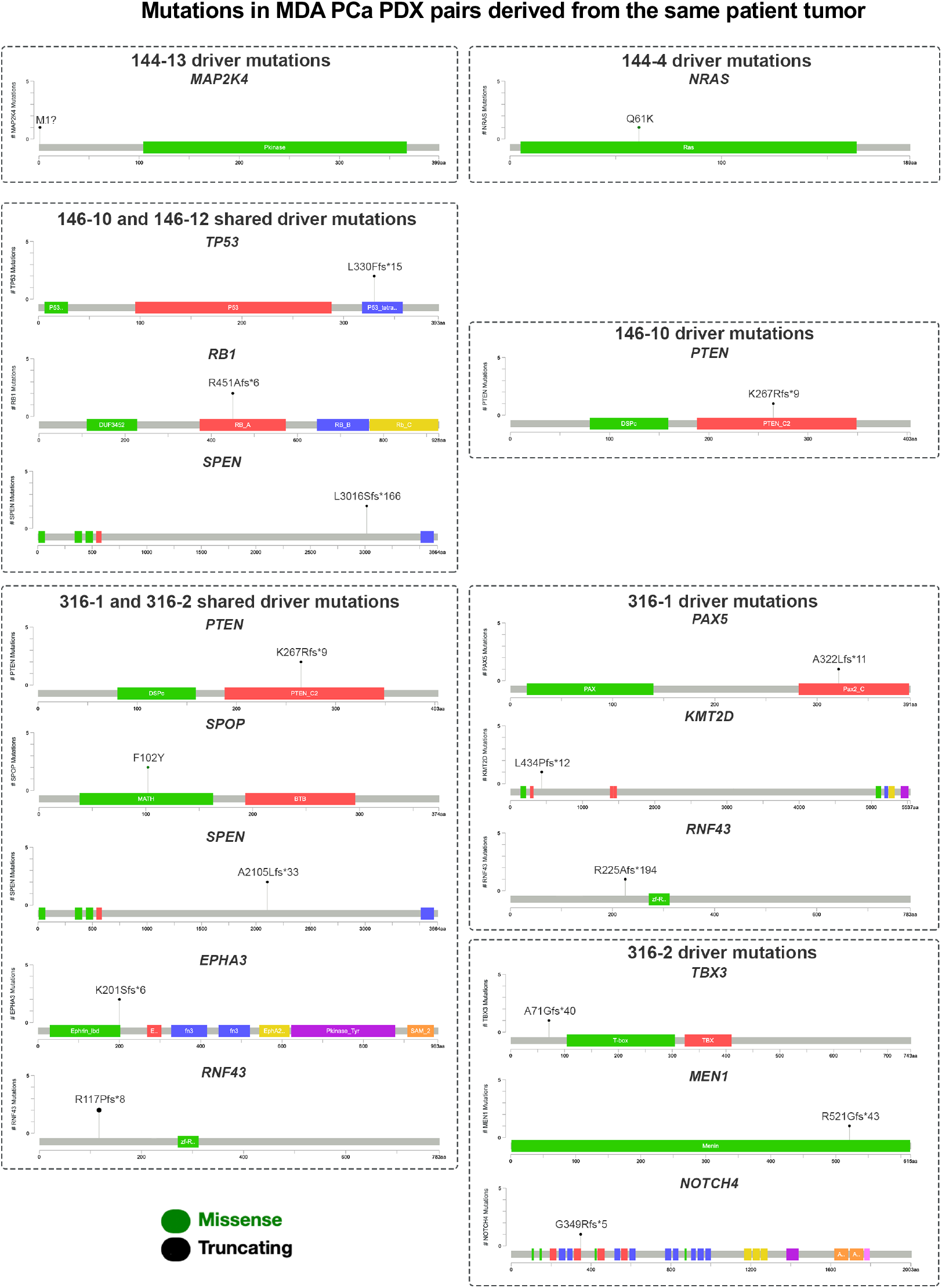
Mutations in MDA PCa PDX pairs derived from the same patient tumor. Lollipop representation of known prostate cancer driver mutations in models derived from two areas of the same tumor. The following pairs are represented: 144-13 and 144-4; 146-10 and 146-12; 316-1 and 316-2. Lollipops were adapted from cBioPortal. For detail on gene domains (represented as color rectangles) refer to cBioPortal.

MDA PCa 146-10 and 146-12 were derived from different areas of a mixed adenocarcinoma and neuroendocrine carcinoma in a primary PCa. They subsequently established as a pure neuroendocrine (MDA PCa 146-10) or an adenocarcinoma (MDA PCa 146-12). Both share mutations in *TP53* (L198Ffs*15) and *RB1* (R451Afs*6), known drivers of PCa (Fig 7, Table 2). Also, they have a mutation in *SPEN* (L3016Wfs*6), a hormone inducible transcription repressor, recently implicated in PCa (25). On the other hand, in addition to their morphological differences, specific genomic alterations were evident in each model. As expected, the neuroendocrine MDA PCa 146-10 does not express AR or its target, ZBTB16 (Table 2). Furthermore, while both PDXs have a heterozygous deletion and low expression of *PTEN*, only MDA PCa 146-10 has a mutation in this gene (K267Rfs*9) (Fig 7, Table 2, Fig S2).

MDA PCa 316-1 was developed from a bone metastasis (adenocarcinoma). MDA PCa 316-2 was developed from CTC obtained at the same time in the blood stream of the same patient. They share mutations in some PCa driver genes: *PTEN* (K267Rfs*9), negative regulator of Wnt pathway *RNF43* (R117Pfs*8) (16) and *SPOP* (F102Y). Also, both have a mutation in the above-mentioned *SPEN* (A2105Lfs*33). However, the bone-derived PDX, MDA PCa 316-1, has a mutation on *PAX5* (A322Lfs*11) (26) and *KMT2D* (L434pfs*12) (27), while the CTC-derived PDX, MDA PCa 316-2, has a mutation in *TBX3* (A71Gfs*40) (28) (Fig 7, Table 2). These findings support the relevance of the environment in the selection of specific characteristics of the cancer cells.

Overall, these results highlight the implications of different pathways promoting progression within the same tumor. These models from different areas of a single tumor are helpful for fully understanding the unique biology of the tumor and depict most accurate treatment strategies.

### Longitudinal studies

We also developed PDXs obtained at different times during the progression of the disease (longitudinal samples), namely MDA PCa 177-B and 189-1 (Patient ID 15), MDA PCa 183-A and 203-A (Patient ID 18), MDA PCa 280-9 and 306-14 (Patient ID 27), and MDA PCa 342-B and 355-9 (Patient ID 37) (Table 3).

**Table 3.**
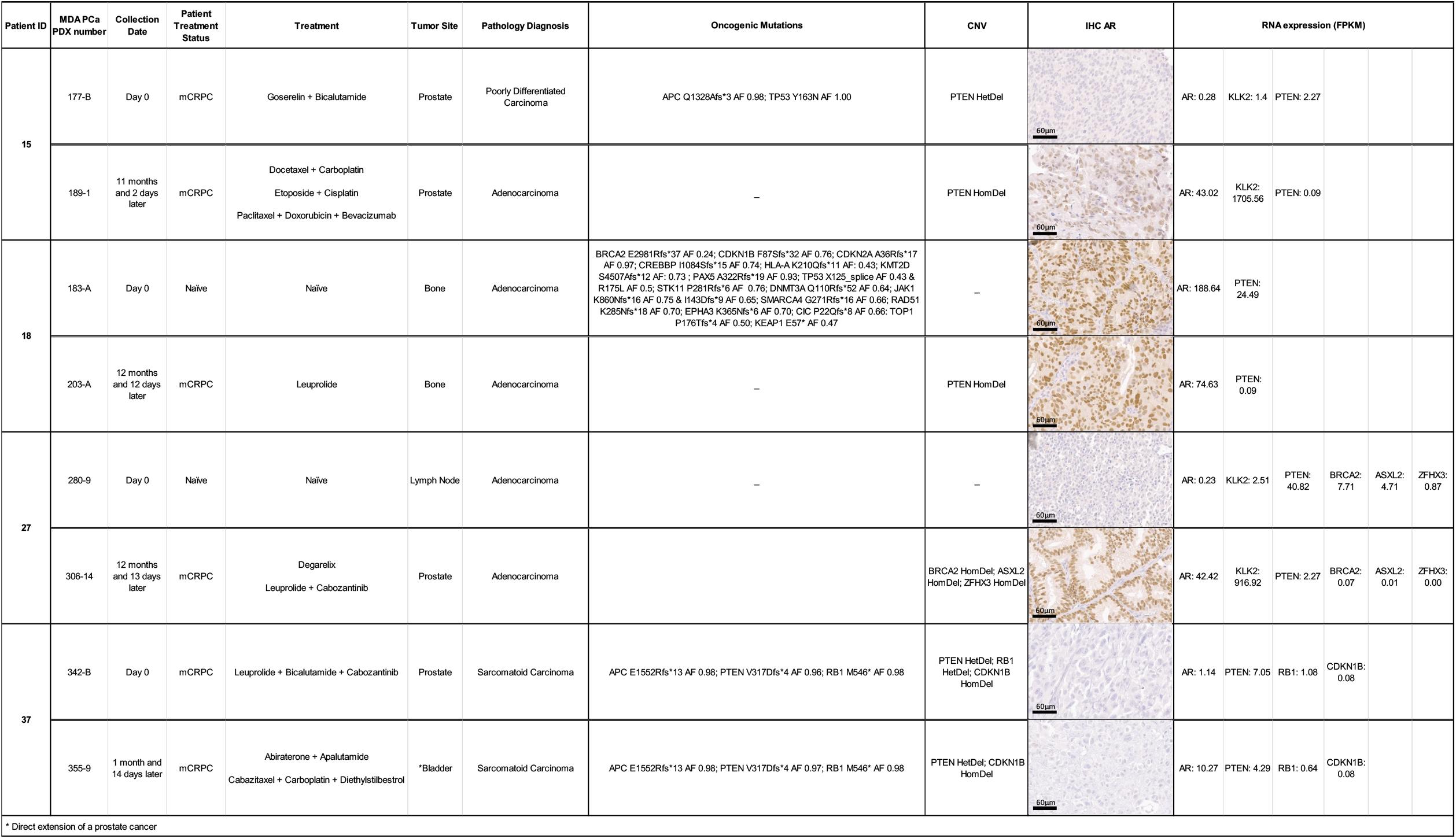
Clinical, morphological and molecular features of longitudinal samples in MPA PCa PDX cohort studied.

In the case of MDA PCa 177-189 pair, both were obtained 11 months apart from the prostate of a man with mCRPC. After the first PDX was developed (MDA PCa 177-B), the patient received a combination of various chemotherapeutics (Table 3). These longitudinal models diverge morphologically, being MDA PCa 177-B characterized as a poorly differentiated carcinoma, with *null* AR expression as assessed by IHC, while MDA PCa 189-1 is an adenocarcinoma that expresses AR. Surprisingly, some molecular alterations detected initially in MDA PCa 177-B (*APC, TP53*) were not found after the second set of treatment. *PTEN* has a heterozygous deletion in MDA PCa 177-B, and MDA PCa 189-1 has a homozygous deletion with loss of expression (Table 3). This suggests that treatment leads to clone selection or homogenization of molecular alteration selecting AR expression and *PTEN* deletion as PCa drivers.

In the other pair (MDA PCa 183-203), MDA PCa 183-A is derived from a bone biopsy of a treatment-naïve adenocarcinoma. The patient was placed on Leuprolide (LHRH agonist), and about 1 year later we obtained a biopsy from a bone metastasis (MDA PCa 203-A). Interestingly, MDA PCa 183-A displayed an abundance of molecular alterations (Table 3); however, the sample obtained after treatment only presented *PTEN* deletion, suggesting homogenization of the tumor after treatment.

The MDA PCa 280-306 pair is a unique case, as MDA PCa 280-9 was obtained from a therapy-naïve lymph node metastasis that does not have any molecular alterations of best known PCa drivers (*PTEN, RB1, TP53, AR*), only AR *null* expression by IHC and low expression by RNAseq (Table 3). The patient was placed on antiandrogen therapy and cabozantinib. About one year later, we developed a PDX (MDA PCa 306-14) derived from the prostate of the same patient. In this case, several mutations were detected (Table 3), suggesting that a lymph node metastasis may not represent a biological meaningful step wise progression.

The MDA PCa 342-355 pair was developed about one month apart from the prostate or local extension to the bladder and they present identical alterations (Table 3), suggesting tumor homogeneity, evidencing that the differences seen in the above models are a consequence of the treatment.

In conclusion, our data suggest that treatment leads to homogenization and/or selection of cells with certain PCa drivers. This warrant further studies.

### Organoids

Historically there has been a limited availability of PCa cell lines that accurately reflect the human disease in its various manifestations (29). In fact, PCa cells typically have poor *in vitro* growth capability as monolayers, and few establish as cell lines; as a consequence, a handful of PCa cell lines representing only a small spectrum of the disease is available.

The recent establishment of *in vitro* growing conditions for PCa cells as organoids (30) is a breakthrough in terms of the possibility of having an *in vitro* methodology to complement PDXs as preclinical models of PCa. However, in our experience and that of others (30), establishing organoids from tumor cells obtained from samples collected after surgery or biopsies from PCa patients has a very low take rate. The majority only grow for a few days to a week.

Therefore, using the organoids technology (30) to grow PCa cells derived from PDXs, we first tested whether this would be an adequate approach to screen for sensitivity to drugs. We started by using widely used PCa cell lines, DU-145, LNCaP, PC3, and VCaP, growing in monolayers and as organoids, and found that the drug responses were very similar between platforms (Fig S3). We then selected some PDX-derived organoids to assess their utility for *in vitro* testing, choosing models derived from naïve adenocarcinoma from bone (MDA PCa 183-A) or from primary (MDA PCa 173-2), and from adrenal gland metastasis of ductal adenocarcinoma (MDA PCa 250-15) (Fig 8). We performed morphologic analysis by H&E (Fig 8A). Next, we tested organoids with chemotherapeutic agents: Cisplatin, Paclitaxel, Cabazitaxel; AR inhibitors: Casodex (Bicalutamide) and Enzalutamide, and PARP inhibitor: Niraparib (Fig 8B). Despite most PDX-derived organoids can grow for few passages, we used the organoids obtained directly from fresh PDX (organoid passage 0) for drug testing, since it is the most accurate approach to standardize the conditions among the models. Of the drugs tested on MDA PCa 183-A, Paclitaxel, Cabazitaxel, Casodex, Enzalutamide and Niraparib reduced cell viability compared with vehicle, with Casodex, Enzalutamide and Niraparib showing a dose-dependent response (Fig 8B, Fig S4). On MDA PCa 173-2, Cabazitaxel, Casodex, Enzalutamide and Niraparib reduced cell viability compared with organoids treated with vehicle, only Casodex showing a dose-dependent response. MDA PCa 250-15 organoids had significant reduced cell viability when treated with Paclitaxel and Casodex in a dose-dependent manner compared with vehicle, but no effect with Cisplatin. These results show that this method could be used as an informative platform for drug screening.

**Figure 8.**
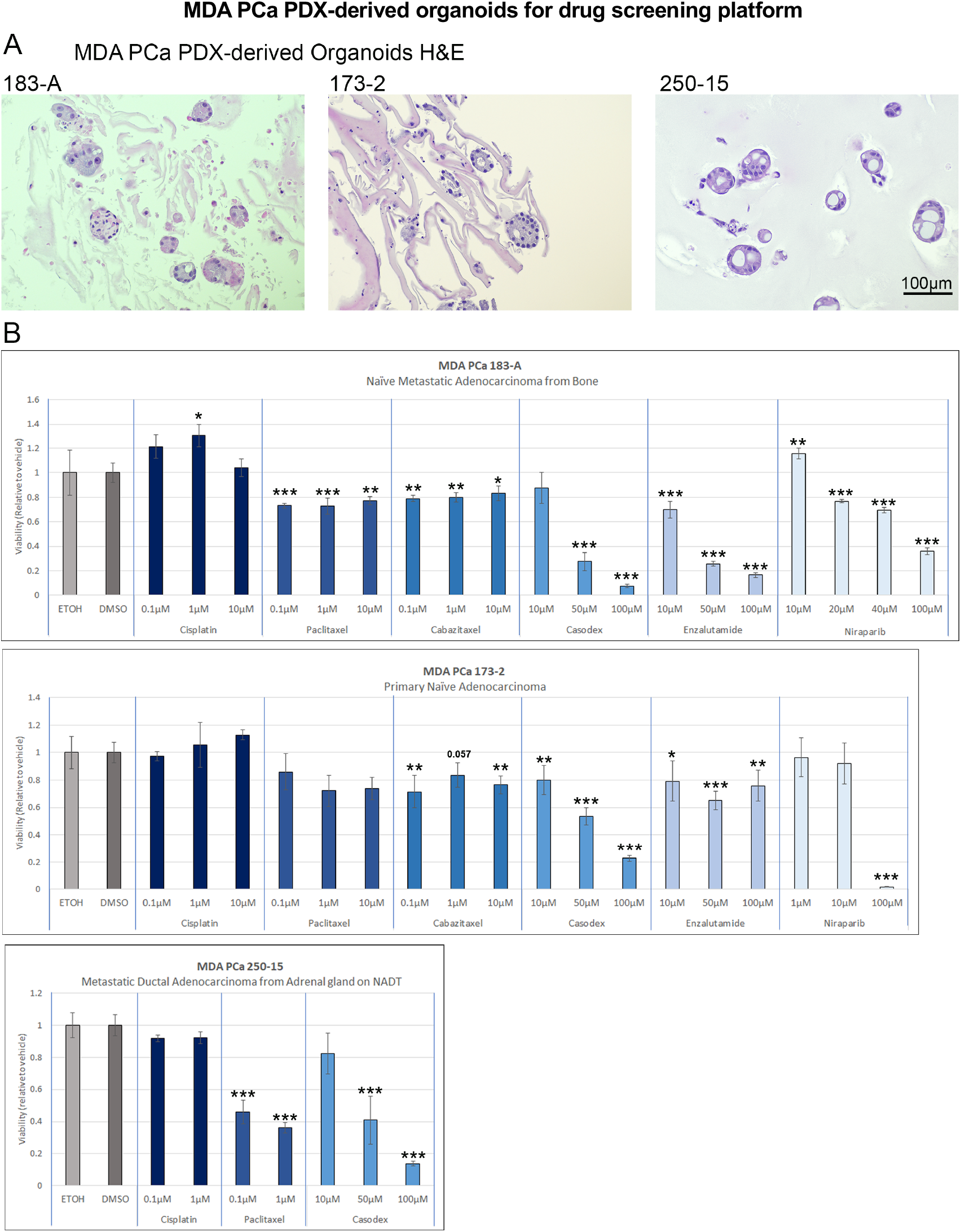
MDA PCa PDX-derived organoids for drug screening platform. **A**. Representative H&E images of organoids derived from MDA PCa PDX **B**. PDX-derived organoids were used to assess the response to different concentrations of drugs (Cisplatin, Paclitaxel, Cabazitaxel, Bicalutamide (Casodex), Enzalutamide, Niraparib) commonly used in the clinic. * P<0.05; ** P<0.01; *** P<0.001. ETOH: ethanol; DMSO: dimethyl sulfoxide; NADT: on neoadjuvant androgen deprivation therapy.

## Discussion

Since patient-derived models of cancer (PDMCs; e.g., PDXs and organoids) were developed, they have led to therapeutically relevant approaches (1-6). The success of these models and recent large-scale genomics studies that have identified deregulated pathways in mCRPC further drives the impetus to understand and improve the utility of PDMCs in addressing clinical gaps that limit progress. Using our expertise applied to critical pathways implicated in CRPC and tissue derivatives of tumor specimens obtained at single time points during PCa progression, organoids technology will complement PDXs and donor human tissue to form *integrated models to study*. Integrated models, composed of human donor tumor and corresponding PDMCs with common molecular alterations, will help prioritize mechanisms of response and resistance to targeted therapy.

In this work, we performed WGS, targeted sequencing, and RNAseq of 46 MDA PCa PDX models derived from 39 patient tumors. The molecular and morphological analyses of each PDX were all performed in representative samples of the same tumor, carefully selected by pathological analysis. This design facilitates the integrated analysis of the different molecular approaches (i.e., WGS, targeted sequencing and RNAseq) with the morphologic and immunoassays results.

We found that heterozygous deletion or amplification of specific genes would not impact on their expression as assessed by RNAseq. However, most homozygous deletions resulted in *null* expression (Fig 2 and see cBioportal).

Further, we found that most pathways frequently altered in PCa were also altered in our cohort and that several known fusions reported in PCa were identified in our PDXs: TMPRSS2-ERG, TMPRSS2-ETV4, SLC45A3-ELK4. These provide evidence that the cohort recapitulates molecular alterations frequent in PCa and that it can be used to ask specific biological questions.

Interestingly, when looking at the most frequently altered genes in PCa (*AR, RB1, TP53* and *PTEN*), only 90% of PDXs presented oncogenic molecular alterations in one or more of these genes. It remains to determine the oncogenic driver in the remaining cases. Among other possible PCa drivers, ARv7, the best characterized and most prevalent AR splice variant, is constitutively active and its implications in PCa have been widely studied (31). We detected ARv7 expression in several models. However, most cases with ARv7 expression have alterations in the known PCa drivers mentioned above. Interestingly, MDA PCa 327-2 express ARv7, in the absence of alterations in *AR, RB1, TP53* and *PTEN*, representing a useful model to understand the function of this variant. Altogether, these analyses emphasize the usefulness of this approach as a resource for the identification of potential novel drivers.

PCa is an increasingly recognized heterogeneous disease (32, 33). The lack of uniform response to agents with different mechanisms of action further demonstrates the biological heterogeneity of these tumors and underscores the urgent need to integrate our knowledge of PCa biology into clinical application. Results of our heterogenous samples highlight the implications of different pathways promoting progression within the same tumor. Moreover, tumor specimens (and corresponding PDMCs) obtained overtime from the same patient will help identify the pathways that drive progression and relapse. Interestingly, the homogenization/reduction in mutational load observed among the longitudinal samples in our cohort suggest that treatment resistance may be a consequence of selection of pre-existent alterations in a subpopulation of cells, highlighting the relevance of understanding tumor heterogeneity of PCa, which may have therapeutic implications.

To address the lack of relevant clinically annotated *in vitro* models of PCa that capture the whole spectrum of the disease, we developed PDX-derived organoids that can recapitulate genetic and morphological features of human PCa. This technology provides a useful platform for *in vitro* drug screening, genetic manipulation of PDX cells and experimentation, reducing times and use of animals.

The results presented in this work serve as a proof-of-principle to the feasibility of the use of organoids for drug testing. Our studies showed some similarities in the results with different drugs, however some differences are observed among particular PDXs tested, stressing the importance of having different models and a unique useful tool as this one to perform rapid and efficient testing. In this way, the ability to use PDXs, organoids and human donor tumor (*integrated models to study*) provides a new way to study the pathogenesis of PCa and accelerate discovery of effective therapies.

## Material and Methods

### Animals

All animal experiments were conducted in accordance with accepted standards of animal care and were approved by the Institutional Animal Care and Use Committee at The University of Texas MD Anderson Cancer Center (Houston, TX).

PDX propagation and harvesting was performed as described (12).

### Selection of MDA PCa PDX

Of our cohort, we selected 46 PDXs derived from 39 men with potentially lethal PCa. The initial selection was based on PDXs with a growth rate that would allow experimentation. The second criteria were based on morphology, namely we selected PDXs that had high tumor content, were homogeneous and had no or minimal necrosis. Each selected PDX tumor was divided in representative samples to performed NGS (including paraffin sections).

### DNA and RNA preparation

Was performed by the Biospecimen Extraction Facility (MD Anderson).

### NGS

We performed WGS, targeted sequencing (T200.1) and RNAseq (Ilumina NGS) in ATGC at MD Anderson Cancer Center. Data was processed at the Department of Genomic Medicine (http://gm.mdanderson.org). T200.1 panel includes 263 genes implicated in the pathogenesis of solid cancers (Table S1).

### Data Processing

NGS data was processed with established in-house bioinformatics pipeline. In brief, raw sequencing base call (BCL) files were first converted into FASTQ files using Illumina bcl2fastq2 conversion software v2.20. DNA sequences were then aligned to the hg19 +mm10 reference genome using the BWA. Picard toolset was used to convert the data into a BAM format with duplicate reads (v1.112, http://broadinstitute.github.io/picard/). Finally, GATK toolkit was used to perform local realignments. Reads mapped to mm10 was filtered to keep HumanOnly (hg19) alignments. MuTect was used to identify somatic single nucleotide polymorphisms (SNPs) and small insertions and deletions (indels). ANNOVAR was applied to annotate each genetic variant with coding sequence change and allele frequency in control populations, including Exome Aggregation Consortium (ExAC), Genome Aggregation Database (gnomAD), the 1000 Genome Project and NHLBI-ESP 6500 exomes.

The following filters were applied to select somatic mutations of good sequencing quality: (1) total reads: ≥ 20 reads in the tumor sample and ≥ 10 reads in normal sample (matched germline sample or merged normal tissues); (2) variant allele frequency (VAF): for SNVs, VAF ≥ 0.02 in tumor sample and ≤ 0.02 in normal sample; for indels, VAF in tumor sample ≥ 0.05 and not observed in normal tissue; (3) length of indels ≤ 100 bp; (4) exclusion of intronic and intergenic mutations, (5) population allele frequency < 0.01 in all of the four control databases.

For the purpose of this publication we focused in known PCa related genes (19, 34, 35); https://cancer.sanger.ac.uk/census)

Somatic CNVs were identified from WGS data in comparing against a common normal consisting of 11 lung blood samples using HMMcopy. The gene-level copy number was assessed by calculating the mean value of the derived log2 scores of tumors versus normal reads for each gene. Copy number status was further categorized using log2.mean cutoffs: >0.4, Amp; −2 ∼ −0.4, HetDel; <-2 HomDel.

Gene fusion events were assessed in both WGS and RNAseq data. For assessing the data comprehensively, seven bioinformatics software were used, including 3 tools on WGS data (brass, delly, and lumpy) and 4 on RNAseq (defuse, mapslice, tophatfusion, fusionmap). Due to a possible discrepancy in breakpoint identification, the findings from different tools were first summarized as gene-pair-level. We used IGV Genome Browser to fine-map the breakpoint.

### ARv7

ARv7 was identified first by RNA reads in IGV. We considered a cutoff of 5 reads for a valid call. We used sashimi plots in IGV to identify putative ARv7 variants. We confirmed the presence of ARv7 by IHC, using standard techniques and Anti-Androgen Receptor (AR-v7 specific) Rabbit Monoclonal Antibody, Clone RM7; RevMab Biosciences (San Francisco, CA). Staining was evaluated by semi-quantitative analysis of pattern, percentage of cells and intensity on a scale of 1 to 3.

### Organoids

PDX-derived organoids were developed from fresh PDXs. Briefly, the tissue was chopped into small pieces; digested with Collagenase II (R&D, Minneapolis, MN) on advanced DMEM/F-12 (gibco, Waltham, MA) with 1% PenStrep (Sigma-Aldrich, St. Louis, MO), 10 mM HEPES (gibco, Waltham, MA) and 1X GlutaMax (gibco, Waltham, MA) on an orbital shaker at 37C for 2 hours; and passed sequentially through a 100 and 70 μm strainer. After centrifugation, cells were incubated with 5 ml of ACK lysis solution (Quality Biological, Gaithersburg, MD) for 5 minutes, when the reaction was stopped with media. Ten thousand cells/well were plated in 20 μl Corning Reduced-Growth Factor Matrigel (Corning, NY) in a 96-well plate. Cells were maintained in PCM media. Drug testing: organoids were treated with Cisplatin, Paclitaxel, Cabazitaxel, Casodex, Enzalutamide and/or Niraparib (Selleckchem, Houston, TX) or vehicle (ethanol [for Cisplatin] or DMSO) for 96h. Viability was evaluated with Promega CellTiter-Glo 3D Cell Viability Assay (Madison, WI), using equivalent volume of reagent. In parallel, 2×10^6^ cells were plated in a dome in 200 μl Reduced-Growth Factor Matrigel in chamber slides. Once organoids developed, they were fixed with 10% formalin ON, transferred to 70% ethanol and paraffin-embedded for histology.

PCM media was prepared as in (36) with the following modifications: 10% (5 ml) Noggin conditioned media and 5% (2.5 ml) R-spondin condition media. *N*-acetyl-L-cysteine and SB202190 were not added. One-way ANOVA was used to assess statistical significance between treatments for each model, followed by Dunnett post-test of each treatment compared with vehicle.

### Statistical analysis

Differences between NE and Ad were assesed with Fisher exact test.

## Acknowledgments

This work was supported by the Prostate Cancer Foundation, NCI Cancer Center Support Grant (P30CA16672), Cancer Center Prostate Cancer SPORE (NIH/NCI P50 CA140388), David H. Koch Center for Applied Research in Genitourinary Cancers at MD Anderson (Houston, TX), and NIH/NCI U01 CA224044

## PDX availability

PDX are available through a material transfer agreement.

