## Supplementary material for "Integrative analysis of the MD Anderson Prostate Cancer Patient-Derived Xenograft Series (MDA PCa PDX)"

**Table S1. T200.1 panel: genes implicated in the pathogenesis of solid cancers**

| | | | | | |
|---|---|---|---|---|---|
| ABL1 | CDKN1B | GABRA6 | MUTYH | RPTOR | XPO1 |
| ABL2 | CDKN2A | GATA1 | MYD88 | RUNX1 | ZRSR2 |
| ACVR1B | CDKN2C | GATA2 | NBN | RUNX1T1 | |
| ACVR2A | CEBPA | GATA3 | NCOR1 | SDHB | |
| AJUBA | CHEK1 | GNA11 | NF1 | SDHC | |
| AKT1 | CHEK2 | GNAQ | NF2 | SDHD | |
| AKT2 | CIC | GNAS | NFE2L2 | SETBP1 | |
| AKT3 | COL2A1 | GSK3B | NKX2-1 | SETD2 | |
| AKTIP | CREBBP | H3F3A | NOTCH1 | SF3B1 | |
| ALK | CSF1R | H3F3B | NOTCH2 | SMAD2 | |
| AMER1 | CTCF | HIST1H3B | NOTCH3 | SMAD3 | |
| APC | CTLA4 | HLA-A | NOTCH4 | SMAD4 | |
| AR | CTNNB1 | HNF1A | NPM1 | SMARCA2 | |
| ARAF | CYLD | HRAS | NRAS | SMARCA4 | |
| ARID1A | CYP2C19 | HSP90AB1 | NSD1 | SMARCB1 | |
| ARID1B | DAXX | IDH1 | NTRK1 | SMARCD1 | |
| ARID2 | DDR2 | IDH2 | NTRK3 | SMC1A | |
| ASXL1 | DDX3X | IGF1R | PALB2 | SMC3 | |
| ATM | DICER1 | IL7R | PAX5 | SMO | |
| ATR | DNMT3A | JAK1 | PBRM1 | SOCS1 | |
| ATRX | EGFR | JAK2 | PDCD1 | SOS1 | |
| AURKA | ELF3 | JAK3 | PDGFRA | SOX9 | |
| AURKB | EP300 | KDM5C | PDGFRB | SPEN | |
| AXIN1 | EPCAM | KDM6A | PHF6 | SPOP | |
| AXIN2 | EPHA2 | KDR | PIK3CA | SRC | |
| AXL | EPHA3 | KEAP1 | PIK3CG | SRSF2 | |
| B2M | EPHA5 | KIT | PIK3R1 | STAG2 | |
| BAP1 | ERBB2 | KMT2A | PLCG1 | STAT3 | |
| BCL11A | ERBB3 | KMT2C | PMS2 | STK11 | |
| BCL2 | ERBB4 | KMT2D | POLE | STK19 | |
| BCOR | ERCC2 | KRAS | PPM1D | SUFU | |
| BIRC2 | ERCC3 | LRP1B | PPP1R3A | SYK | |
| BRAF | ERCC4 | MAP2K1 | PPP2R1A | TBC1D4 | |
| BRCA1 | ERCC5 | MAP2K2 | PRDM1 | TBX3 | |
| BRCA2 | ESR1 | MAP2K4 | PREX2 | TERTp | |
| BTK | ETV1 | MAP3K1 | PRG4 | TET2 | |
| CARD11 | EZH2 | MAP3K13 | PTCH1 | TGFB1 | |
| CASP8 | FADD | MAP3K4 | PTEN | TGFBR1 | |
| CBL | FANCA | MAPK1 | PTK2 | TGFBR2 | |
| CCND1 | FANCD2 | MCL1 | PTPN11 | TNF | |
| CCND2 | FBXW7 | MDM2 | PTPRB | TNFAIP3 | |
| CCND3 | FGFR1 | MED12 | RAC1 | TOP1 | |
| CCNE1 | FGFR2 | MEN1 | RAD51 | TOP2A | |
| CD274 | FGFR3 | MET | RAD51C | TP53 | |
| CD79A | FGFR4 | MITF | RAF1 | TSC1 | |
| CD79B | FH | MLH1 | RARA | TSC2 | |
| CDC27 | FLT1 | MPL | RB1 | TSHR | |
| CDC73 | FLT3 | MSH2 | RET | U2AF1 | |
| CDH1 | FLT4 | MSH6 | RICTOR | VEGFA | |
| CDK12 | FOXA1 | MST1 | RNF43 | VHL | |
| CDK4 | FOXL2 | MST1R | ROS1 | WHSC1L1 | |
| CDK6 | FTO | MTOR | RPS6KB1 | WT1 | |

#### Fig. S1. Mutational landscape of prostate cancer frequently altered pathways

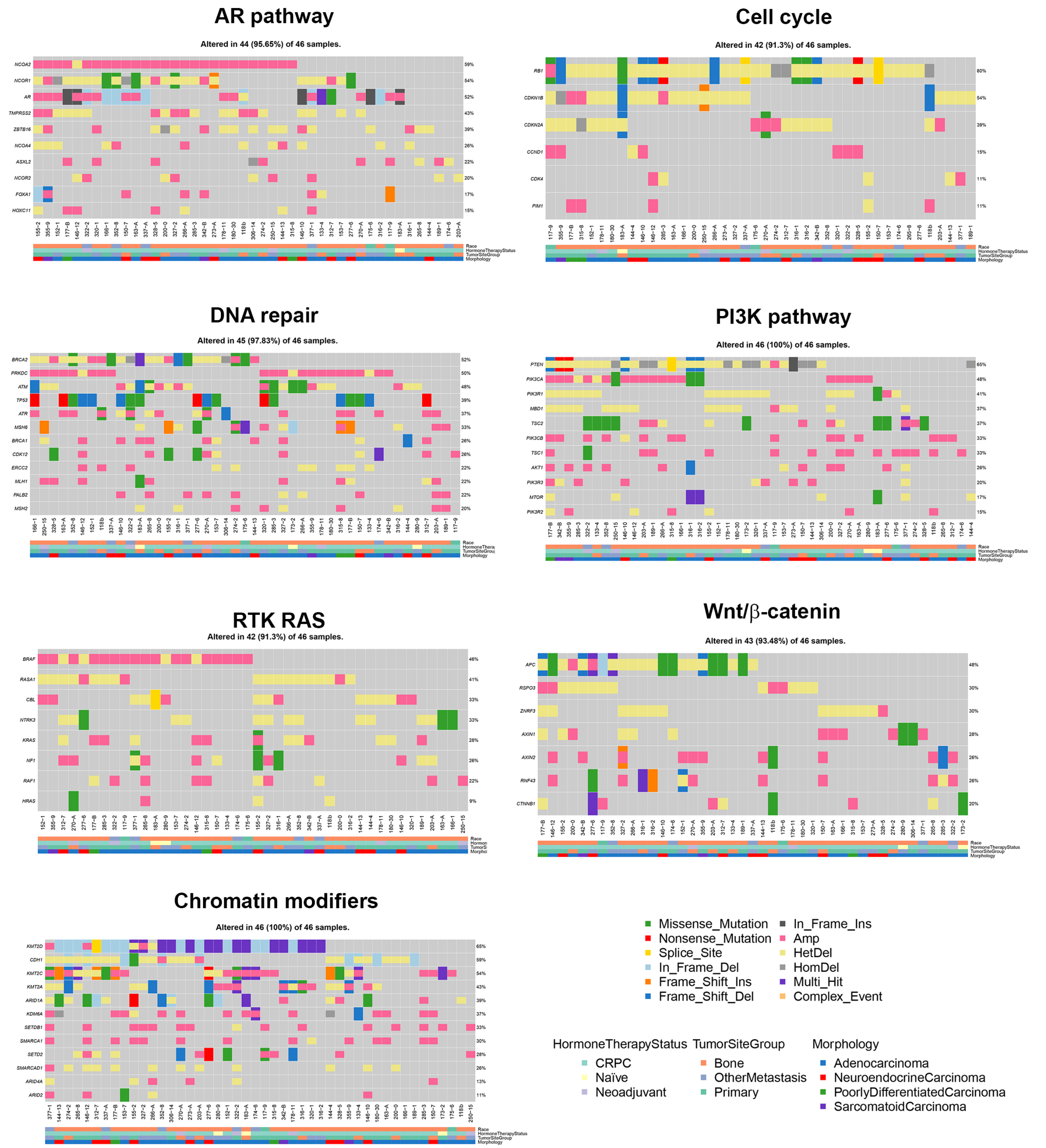


**Figure S1**. Detail of SNP/Indel and CNV identified by T200 and WGS for AR, Cell cycle, DNA repair, RTK-RAS, PI3K, Wnt/β-catenin and Chromatin modifiers pathways.

### Table S2. Fusions identified in 46 MDA PCa PDXs

| Gene.A | Gene.B | PDX | Data Source | Detection meth. | Chr.A | Pos.A | Chr.B | Pos.B | dir.A.B |
|---|---|---|---|---|---|---|---|---|---|
| TMPRSS2 | ETV4 | 175-6 | WGS | bs.dl.lp | chr21 | 42867729 | chr17 | 41632042 | -+ |
| | | | RNAseq | df.fm.mp.th | chr21 | 42867729 | chr17 | 41632042 | -+ |
| | | 183-A | WGS | bs.dl.lp | chr21 | 42866568 | chr17 | 41650089 | -+ |
| | | | RNAseq | df.fm.mp.th | chr21 | 42866568 | chr17 | 41650088 | -+ |
| TMPRSS2 | ERG | 173-2 | WGS | bs.lp | chr21 | 42872537 | chr21 | 39863885 | -+ |
| | | | RNAseq | df.fm.mp | chr21 | 42879877 | chr21 | 39817544 | -+ |
| | | 189-1 | WGS | bs.lp | chr21 | 42860881 | chr21 | 39871051 | -+ |
| | | | RNAseq | df.fm.mp | chr21 | 42861434 | chr21 | 39817544 | -+ |
| | | 203-A | WGS | bs.lp | chr21 | 42860880 | chr21 | 39871050 | -+ |
| | | | RNAseq | df.fm.mp | chr21 | 42861434 | chr21 | 39817544 | -+ |
| | | 133-4 | WGS | bs.lp | chr21 | 42872545 | chr21 | 39860834 | -+ |
| | | | RNAseq | df.fm.mp | chr21 | 42879877 | chr21 | 39817544 | -+ |
| | | 285-3 | WGS | bs.lp | chr21 | 42875598 | chr21 | 39900360 | -+ |
| | | | RNAseq | df.fm.mp | chr21 | 42880008 | chr21 | 39817544 | -+ |
| | | 306-14 | WGS | bs.lp | chr21 | 42859305 | chr21 | 39871943 | -+ |
| | | | RNAseq | df.fm.mp | chr21 | 42860321 | chr21 | 39775631 | -+ |
| | | 152-1 | WGS | bs.lp | chr21 | 42874174 | chr21 | 39861279 | -+ |
| | | | RNAseq | df.fm.mp | chr21 | 42879877 | chr21 | 39817544 | -+ |
| | | 150-7 | WGS | bs.lp | chr21 | 42873453 | chr21 | 39862825 | -+ |
| | | | RNAseq | df.mp | chr21 | 42848504 | chr21 | 39865501 | -- |
| | | 144-4 | WGS | bs.lp | chr21 | 42872479 | chr21 | 39884147 | -+ |
| | | | RNAseq | fm | chr21 | 42879877 | chr21 | 39817544 | -+ |
| | | 320-1 | WGS | bs.lp | chr21 | 42867291 | chr21 | 39821458 | -+ |
| | | | RNAseq | df.fm.mp | chr21 | 42870046 | chr21 | 39817544 | -+ |
| | | 153-7 | WGS | bs.lp | chr21 | 42873029 | chr21 | 39859982 | -+ |
| | | | RNAseq | df.fm.mp | chr21 | 42879877 | chr21 | 39817544 | -+ |
| | | 163-A | WGS | bs.lp | chr21 | 42870627 | chr21 | 39974760 | -+ |
| | | | RNAseq | df.fm.mp | chr21 | 42879877 | chr21 | 39956869 | -+ |
| | | 166-1 | WGS | bs.lp | chr21 | 42870627 | chr21 | 39974760 | -+ |
| | | | RNAseq | df.fm.mp | chr21 | 42879877 | chr21 | 39956869 | -+ |
| FOXA1 | ETV1 | 250-15 | WGS | dl | chr14 | 38060104 | chr7 | 13973676 | -- |
| | | | RNAseq | df.mp | chr14 | 38060113 | chr7 | 13973666 | -- |
| ETV1 | FOXA1 | 250-15 | WGS | bs.lp | chr7 | 13973676 | chr14 | 38060114 | -- |
| | | | RNAseq | th | chr7 | 13974515 | chr14 | 38060606 | -- |
| ACSL3 | ETV1 | 177-B | WGS | bs.dl.lp | chr2 | 223750008 | chr7 | 13975069 | ++ |
| | | | RNAseq | df.fm.mp.th | chr2 | 223725976 | chr7 | 13971374 | ++ |
| ETV1 | ACSL3 | 177-B | WGS | bs.dl | chr7 | 13975168 | chr2 | 223750078 | -- |
| | | | RNAseq | fm.mp | chr8 | 13975333 | chr3 | 223752548 | -- |
| SLC45A3 | ELK4 | 117-9 | WGS | | | | | | |
| | | | RNAseq | df | chr1 | 205630989 | chr1 | 205593019 | -+ |
| | | 153-7 | WGS | | | | | | |
| | | | RNAseq | df | chr1 | 205628617 | chr1 | 205593019 | -+ |
| | | 166-1 | WGS | | | | | | |
| | | | RNAseq | df | chr1 | 205630989 | chr1 | 205593019 | -+ |
| | | 174-6 | WGS | | | | | | |
| | | | RNAseq | df | chr1 | 205628617 | chr1 | 205593019 | -+ |
| | | 175-6 | WGS | | | | | | |
| | | | RNAseq | df | chr1 | 205630989 | chr1 | 205593019 | -+ |
| | | 178-11 | WGS | | | | | | |
| | | | RNAseq | df | chr1 | 205630989 | chr1 | 205593019 | -+ |
| | | 189-1 | WGS | | | | | | |
| | | | RNAseq | df | chr1 | 205628617 | chr1 | 205593019 | -+ |
| | | 200-1 | WGS | | | | | | |
| | | | RNAseq | df | chr1 | 205630989 | chr1 | 205593019 | -+ |
| | | 203-A | WGS | | | | | | |
| | | | RNAseq | df | chr1 | 205628617 | chr1 | 205593019 | -+ |
| | | 306-14 | WGS | | | | | | |
| | | | RNAseq | df | chr1 | 205628506 | chr1 | 205593072 | -+ |
| | | 250-15 | WGS | | | | | | |
| | | | RNAseq | df | chr1 | 205628617 | chr1 | 205593019 | -+ |
| | | 146-12 | WGS | | | | | | |
| | | | RNAseq | df | chr1 | 205630989 | chr1 | 205593019 | -+ |

bs: brass, dl: delly, lp: lumpy, df: defuse, mp: mapslice, th: tophatfusion, fm: fusionmap

#### Fig. S2. CNV in PDX pairs derived from the same tumor

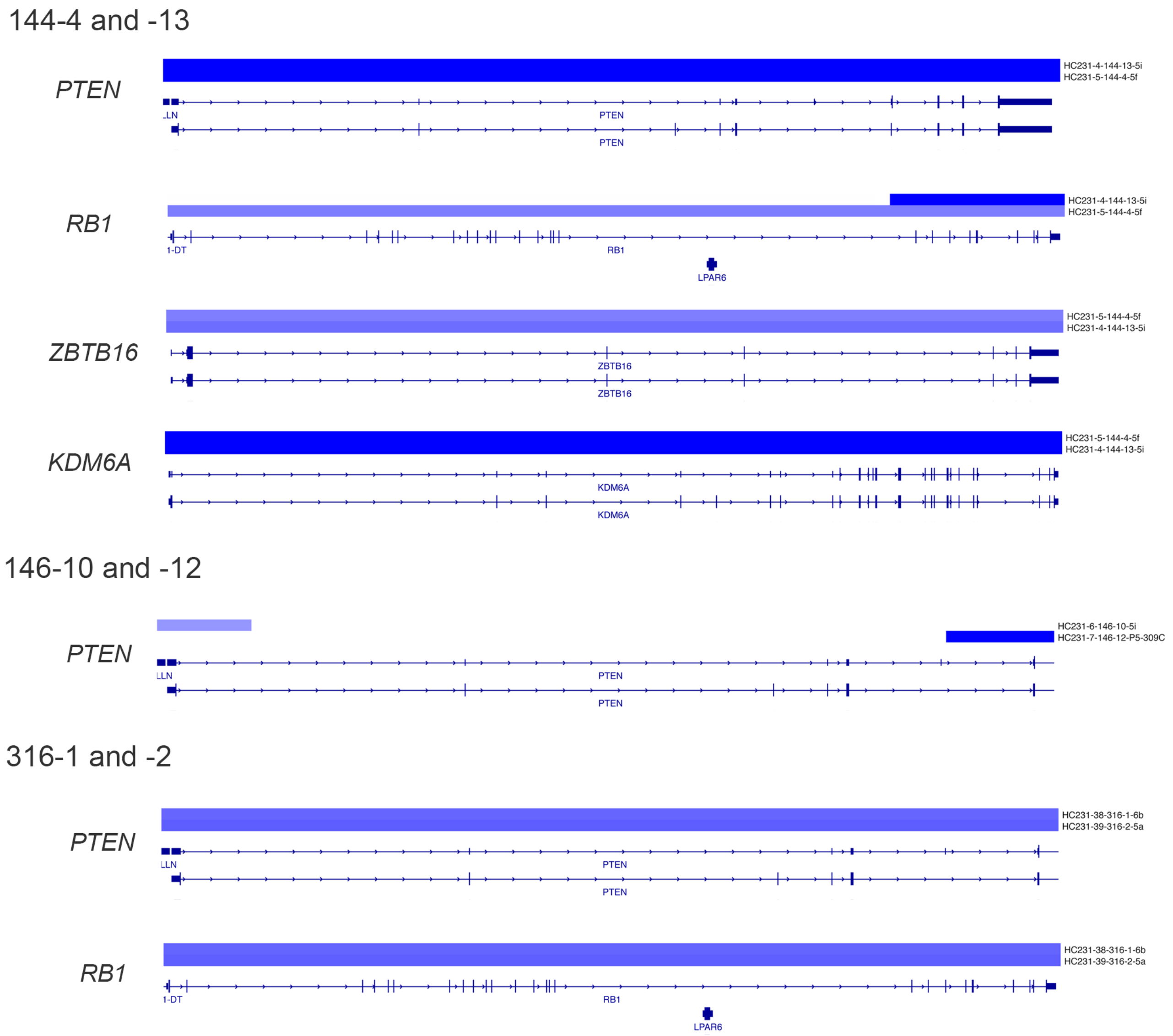


**Figure S2**. CNV based on WGS data of prostate cancer genes in models derived from two areas of the same tumor.

#### Fig. S3. Drug screening on PCa cell lines monolayers and organoids

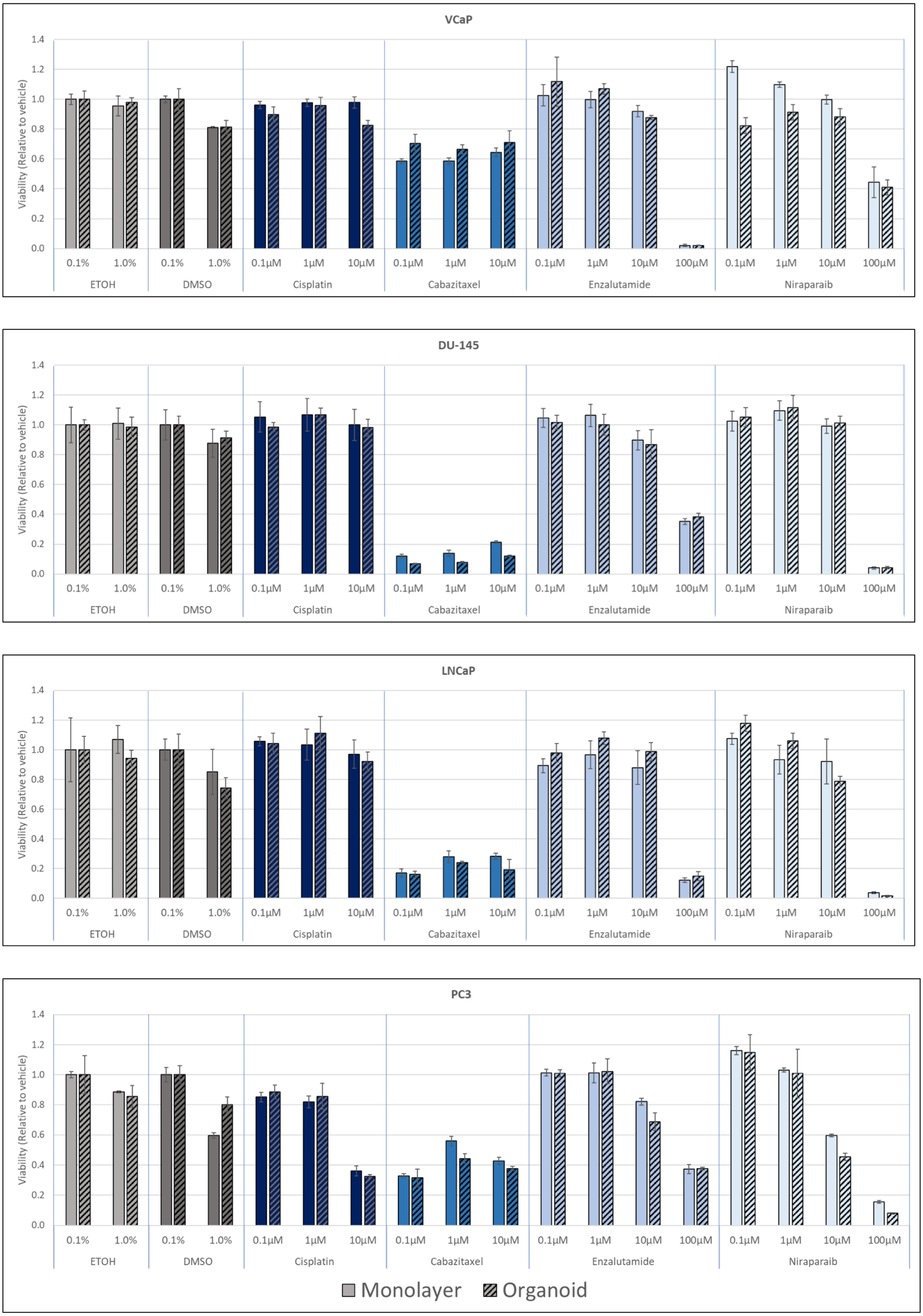


**Figure S3**. PCa cell lines grown as monolayers and organoids were used to assess their response to different concentrations of different drugs commonly used in the clinic. * P<0.05; ** P<0.01; *** P<0.001

**Fig. S4. Drug screening on MDA PCA 183-A derived organoids**

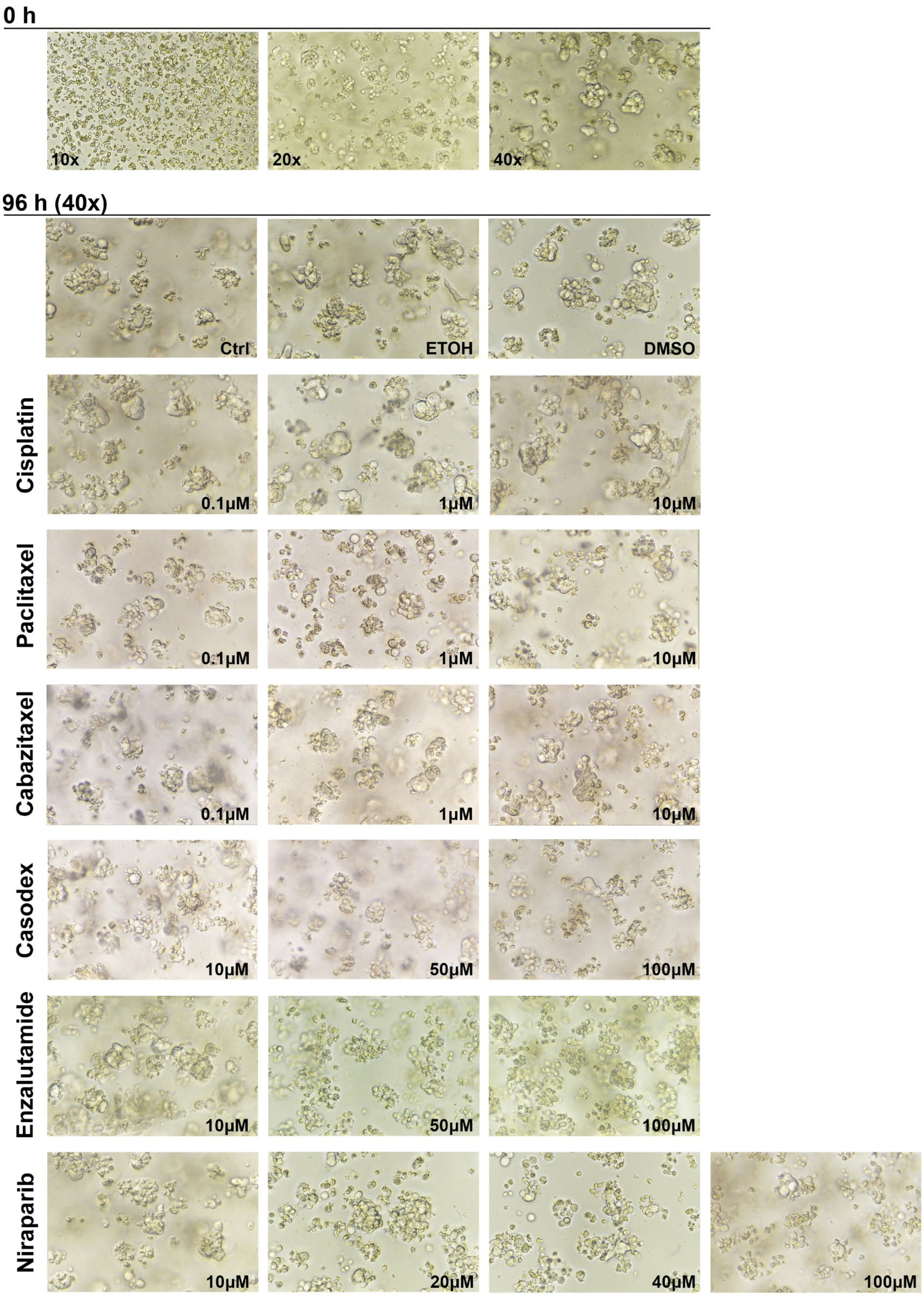


**Figure S4**. Representative Z-stack bright-field images of MDA PCa 183-A derived organoids before (0h) and after (96h) treatment with different concentrations of different drugs commonly used in the clinic or vehicle and control.
